# Complex fitness landscape shapes variation in a hyperpolymorphic species

**DOI:** 10.1101/2021.10.10.463656

**Authors:** A. V. Stolyarova, T. V. Neretina, E. A. Zvyagina, A. V. Fedotova, A. S. Kondrashov, G. A. Bazykin

## Abstract

It is natural to assume that patterns of genetic variation in hyperpolymorphic species can reveal large-scale properties of the fitness landscape that are hard to detect by studying species with ordinary levels of genetic variation^1,2^. Here, we study such patterns in a fungus *Schizophyllum commune*, the most polymorphic species known^3^. Throughout the genome, short-range linkage disequilibrium caused by attraction of rare alleles is higher between pairs of nonsynonymous than of synonymous sites. This effect is especially pronounced for pairs of sites that are located within the same gene, especially if a large fraction of the gene is covered by haploblocks, genome segments where the gene pool consists of two highly divergent haplotypes, which is a signature of balancing selection. Haploblocks are usually shorter than 1000 nucleotides, and collectively cover about 10% of the *S. commune* genome. LD tends to be substantially higher for pairs of nonsynonymous sites encoding amino acids that interact within the protein. There is a substantial correlation between LDs at the same pairs of nonsynonymous sites in the USA and the Russian populations. These patterns indicate that selection in *S. commune* involves positive epistasis due to compensatory interactions between nonsynonymous alleles. When less polymorphic species are studied, analogous patterns can be detected only through interspecific comparisons.

---

Alleles do not affect fitness and other phenotypic traits independently and, instead, engage in epistatic interactions^4–11^. Epistasis is pervasive at the scale of between-species differences, where it is saliently manifested by Dobzhansky-Muller incompatibilities and results in low fitness of interspecific hybrids^12–14^. By contrast, at the scale of within-population variation, the importance of epistasis remains controversial^15–19^. This may look like a paradox, because such variation provides an opportunity to detect epistasis through linkage disequilibrium (LD), non-random associations between alleles at different loci^20–24^. Indeed, epistatic selection generates LD which can be detected^24–28^. Perhaps, the fitness landscape is complex macroscopically but is more smooth microscopically or, in other words, epistasis is genuinely more pronounced at a macroscopic scale^29^. If so, studying epistasis in hyperpolymorphic populations, where differences between genotypes can be as high as those between genomes of species from different genera or even families, holds a great promise because variation within such a population can cover multiple fitness peaks or a sizeable chunk of a curved ridge of high fitness^14,30–33 34,35^ (Supplementary Note 1).

## Elevated LD at nonsynonymous sites

In a vast majority of species, nucleotide diversity π, the evolutionary distance between a pair of randomly chosen genotypes, is, at selectively neutral sites, of the order of 0.001 (as in *Homo sapiens*) or 0.01 (as in *Drosophila melanogaster*)^1^. Still, a few hyperpolymorphic species with π > 0.1 are known, of which the wood-decaying fungus *Schizophyllum commune* is the most extreme, where π = 0.20 or 0.13 in the USA or the Russian populations, respectively^3^ (Supplementary Fig. 1). The two populations of *S. commune* are highly divergent (dS between populations ≈ 0.34, F_st_ = 0.58), but there is essentially no structure within either of them (Supplementary Fig. 2). We studied 34 haploid genotypes from the USA and 21 from Russia and compared the LD between nonsynonymous SNPs (LD_nonsyn_) to that between synonymous SNPs (LD_syn_).

At sites with minor allele frequency (MAF) > 0.05, in both *S. commune* populations, LD_nonsyn_ is much higher than LD_syn_ at the same nucleotide distance (Fig. 1a, Supplementary Fig. 3a). This excess of LD_nonsyn_ is much stronger for pairs of SNPs located within the same gene, compared to pairs of SNPs from adjacent genes at the same distance. By contrast, the excess of LD_nonsyn_ is independent of whether the two SNPs are located within the same or in different exons of a gene (Supplementary Fig. 4). In *S. commune*, the recombination rate is higher within exons^36^, which may affect the patterns of LD; however, this factor could only reduce within-gene LD, and in any case cannot explain the difference between LD_nonsyn_ and LD_syn_.

**Figure 1.**
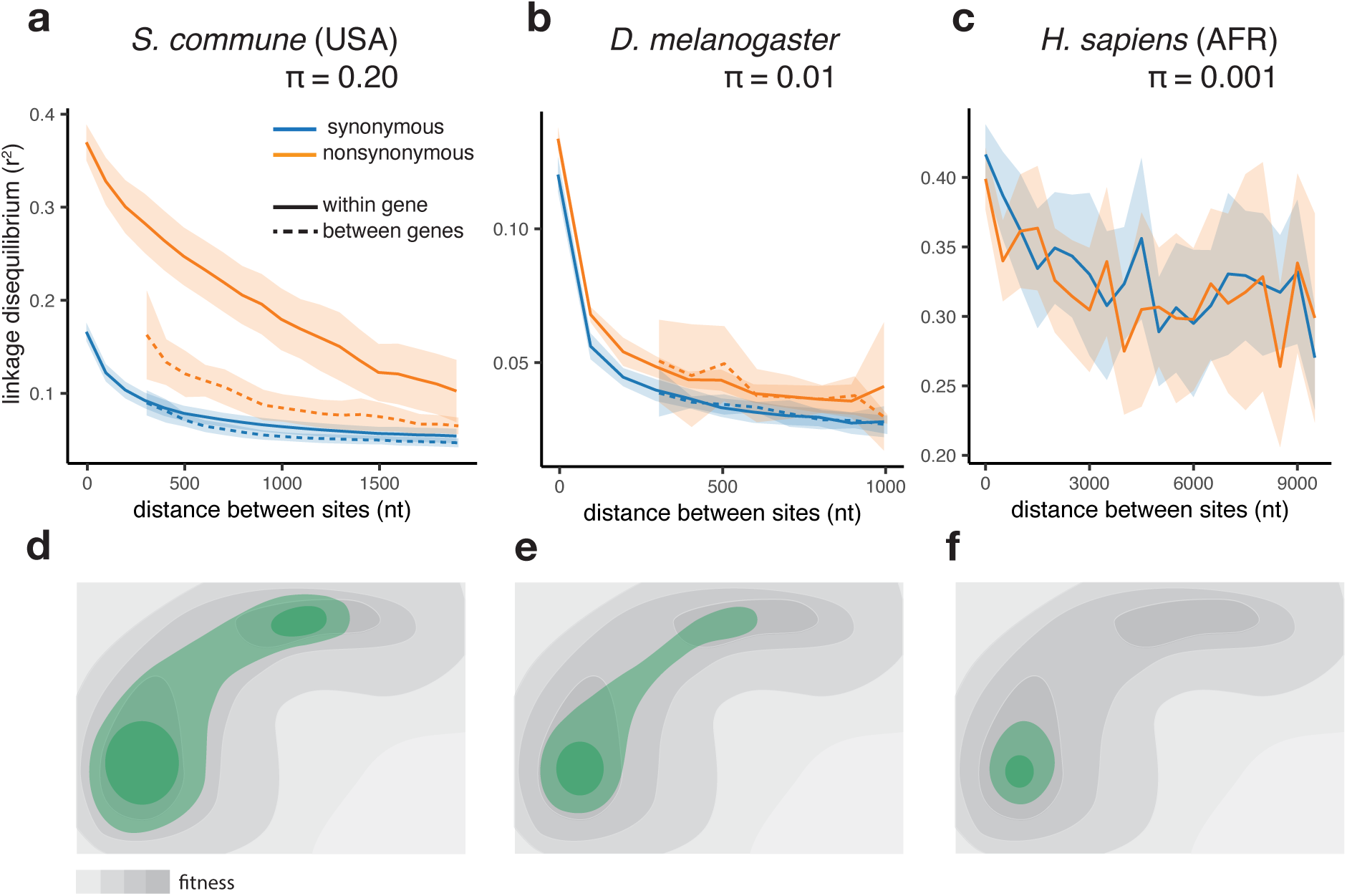
The efficiency of epistatic selection in populations with different levels of genetic diversity. (**a**-**c**) LD in natural populations for SNPs with MAF > 0.05. (**a**) USA population of *S. commune*, (**b**) Zambian population of *D. melanogaster*, (**c**) African superpopulation of *H. sapiens*. Filled areas in (**a**)-(**c**) indicate SE of LD calculated for each chromosome or scaffold separately. (**d-f**) A hyperpolymorphic population (**d**) may occupy a sizeable chunk of a complex fitness landscape, leading to pervasive positive epistasis, while variation within less polymorphic populations (**e** and **f**) is confined to smaller, and approximately linear, portions of the landscape, so that no strong epistasis and LD can emerge. The area of the landscape covered by the population is shown in green.

A much weaker excess of LD_nonsyn_ over LD_syn_ for MAF > 0.05 is also observed in the less genetically diverse *D. melanogaste*r population (Fig. 1b). In the still less polymorphic human populations, LD_nonsyn_ is indistinguishable from LD_syn_ at the same distances (Fig. 1c, Supplementary Fig. 3b).

The excess of LD_nonsyn_ over LD_syn_ corresponds to the attraction between rare alleles at nonsynonymous sites. This attraction can only appear due to positive epistasis between such alleles - higher-than-expected fitness of their combinations (Supplementary Note 2). Positive epistasis can be expected to cause stronger LD in more polymorphic populations (Fig. 1d-f, Supplementary Note 1) and must be more common for pairs of sites located within the same gene, which are more likely to interact with each other. For SNPs with MAF < 0.05, LD_nonsyn_ is similar or lower to LD_syn_ for all the three species, consistent with the effects of random drift, Hill-Robertson interference, and/or negative epistasis (Supplementary Note 3).

## Elevated LD between interacting sites

Natural selection acting on physically interacting amino acids that are located close to each other within the three-dimensional structure of a protein is characterized by strong epistasis which leads to their coevolution at the level of between-species differences^37–39^. Extraordinary diversity of *S. commune* makes it possible to observe an analogous phenomenon at the level of within-population variation. In both *S. commune* populations, pairs of nonsynonymous SNPs are in stronger LD when they are located at codons encoding physically close (within 10Å) than distant amino acids (Fig. 2a; permutation test p-value < 1e-3). This is not the case for pairs of synonymous SNPs (Fig. 2a; permutation test p-value = 0.58).

**Figure 2.**
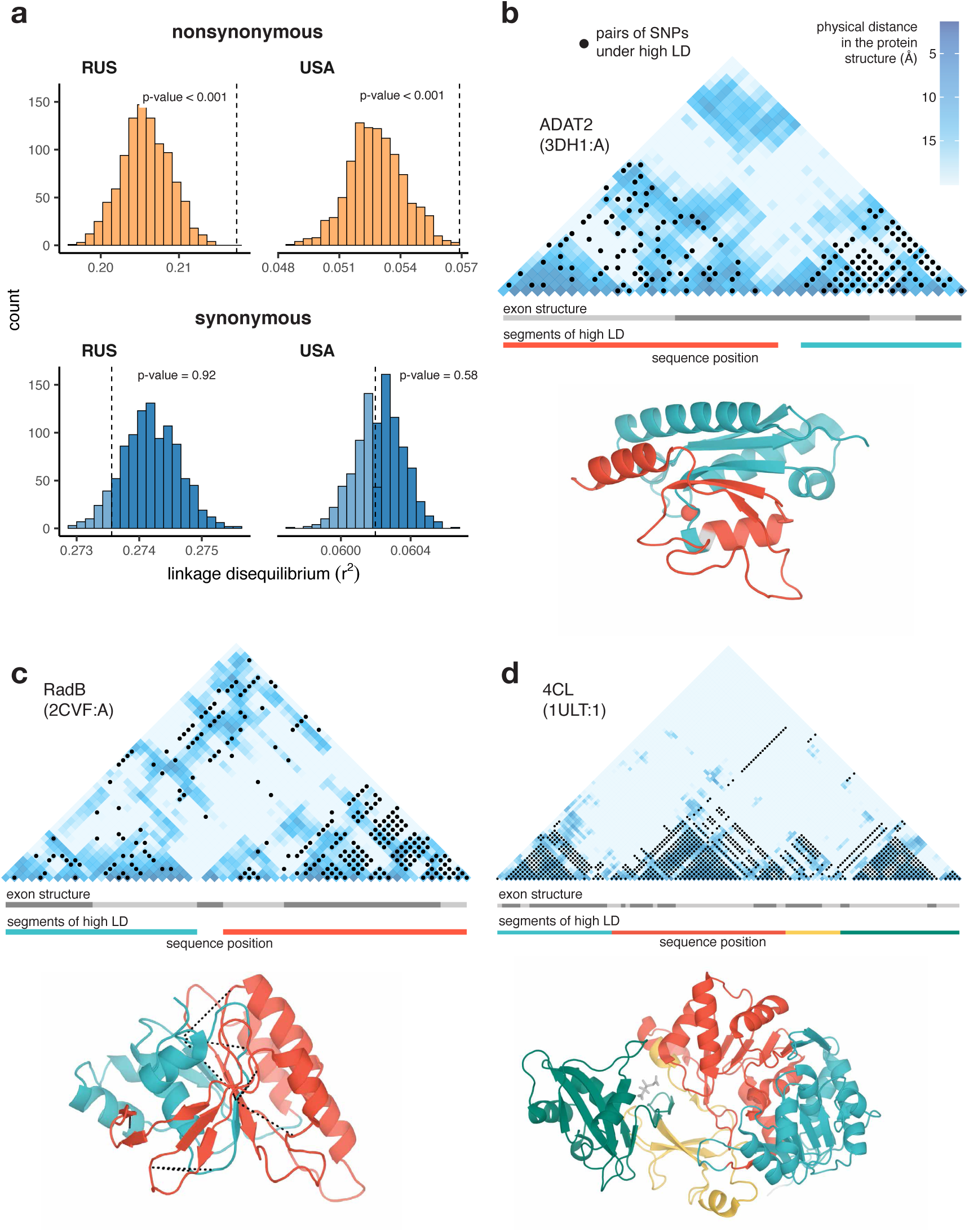
Excessive LD between physically interacting protein sites. (**a**) Within pairs of SNPs that correspond to pairs of amino acids that are colocalized within 10Å in the protein structure, the LD is elevated between nonsynonymous, but not between synonymous, sites. Dashed lines show the average LD. Permutations were performed by randomly sampling pairs of non-interacting SNPs while controlling for genetic distance between them, measured in amino acids; pairs of SNPs closer than 5 aa were excluded. (**b**-**d**) Examples of proteins with LD patterns matching their three-dimensional structures. Heatmaps show the physical distance between pairs of sites in the protein structure; only positions carrying biallelic SNPs are shown. Black dots correspond to pairs of sites with high LD (> 0.9 quantile for the gene). Dashed lines in (**c**) structure show high LD between physically close SNPs from different segments of high LD. In these examples, LD is calculated in the Russian population of *S. commune*.

Moreover, it is possible to identify individual proteins with significant associations between the patterns of LD and of physical interactions between sites. At a 5% FDR, we found 22 such proteins in the USA population, and 87 proteins in the Russian population (Supplementary Table 1); three examples are shown in Fig. 2b-d (see also Supplementary Fig. 5 and 6). The alignment of ADAT2 protein contains two segments (teal and red in Fig. 2b) characterized by high within-segment LD. The boundaries of these segments match those of structural units of the protein, but not the exon structure of its gene. In RadB protein, a similar pattern is observed, and LD is also elevated between pairs of SNPs from different segments on the interface of the corresponding structural units (Fig. 2c). The alignment of 4CL protein can be naturally split into four high-LD segments, which also match its structure (Fig. 2d).

## Distinct regions of high LD

The magnitude of LD varies widely along the *S. commune* genome. Visual inspection of the data shows a salient pattern of regions of relatively low LD, alternating with mostly short regions of high LD (haploblocks, Supplementary Fig. 7). We calculated LD along the genome in a sliding window of 250 nucleotides and regarded as a haploblock any continuous genomic region with LD values that belong to the heavy tail of its distribution (see Methods).

In the USA population, 8.4% of the genome is occupied by 5316 such haploblocks, 56% consist of regions with background LD level, and the rest cannot be analyzed due to poor alignment quality or low SNP density. 88% of the haploblocks are shorter than 1000 nucleotides, although the longest haploblocks spread for several thousand nucleotides (Supplementary Fig. 8). In the Russian population, there are 10694 haploblocks, occupying 15.9% of the genome, and regions of background LD cover 39% of it. There is only a modest correlation between the USA and Russian haploblocks: the probability that a genomic position belongs to a haploblock in both populations is 2.3% instead of the expected 1.3%, indicating their relatively short persistence time in the populations (examples shown in Supplementary Figure 7).

LD within a haploblock is usually so high that most genotypes can be attributed to one of just two distinct haplotypes, which carry different sets of alleles (Supplementary Fig. 9). This results in a bimodal distribution of the fraction of minor alleles in a genotype within a haploblock, because some genotypes belong to the major haplotype and, thus, carry only a small fraction of minor alleles, and other genotypes belong to the minor haplotype and, thus, possess a high fraction of minor alleles (Fig. 3a). Polymorphic sites within haploblocks are characterized by higher MAF than that at sites that reside in non-haploblock regions (t-test p-value < 2e-16), and in the USA population MAFs within a haploblock are positively correlated with its strength of LD (Fig. 3b, Pearson correlation estimate = 0.07, p-value < 2e-6).

**Figure 3.**
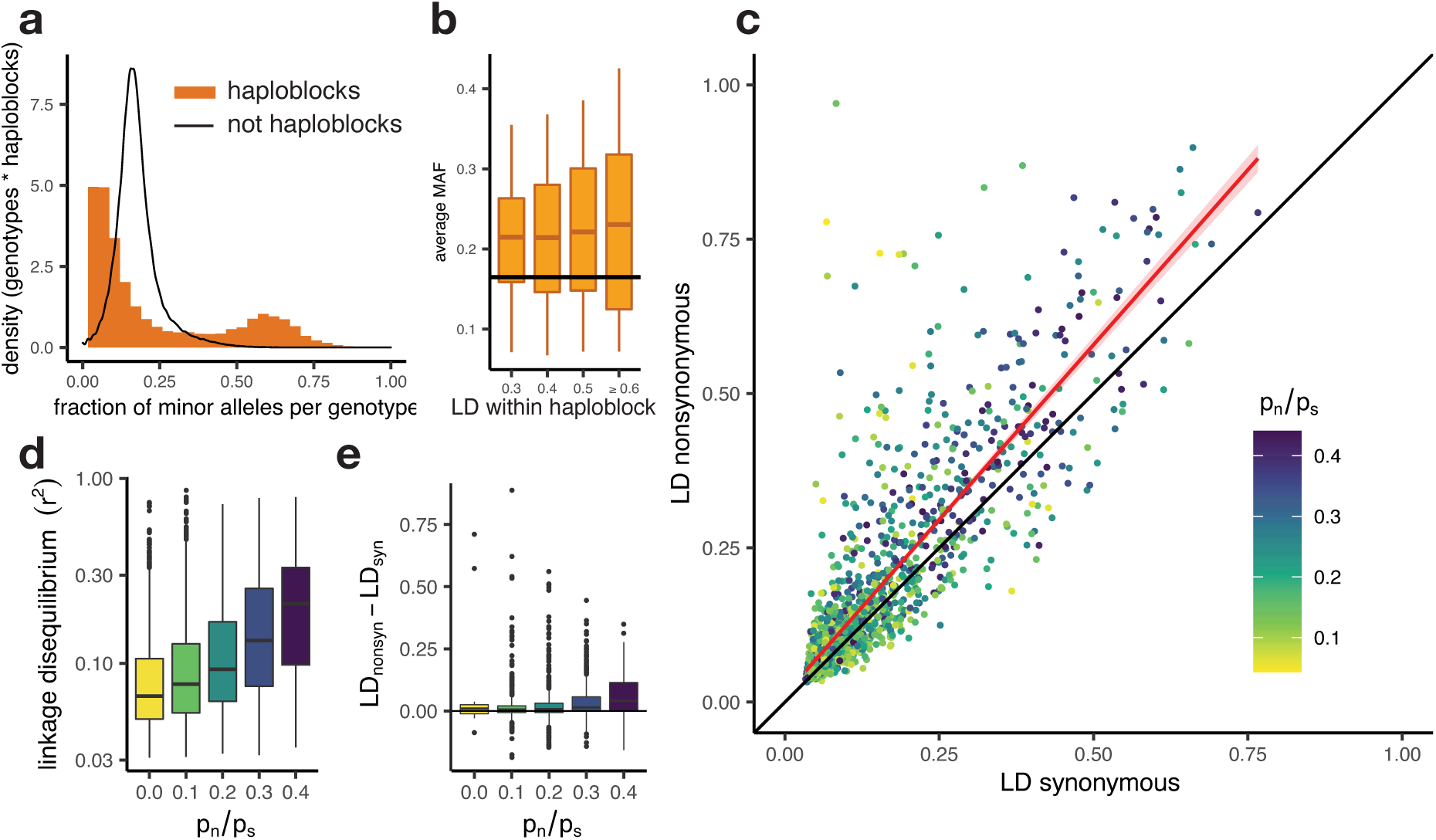
Patterns of linkage disequilibrium in the USA population of *S. commune*. (**a**) Distribution of the fraction of polymorphic sites that carry minor alleles in a genotype within haploblocks. Black line shows the distribution of fraction of minor alleles in genotypes in non-haploblock regions. (**b**) Distributions of the average MAF within a haploblock for haploblocks with different average values of LD. The average MAF in non-haploblock regions is shown as a horizontal black line for comparison. (**c**) LD between nonsynonymous and synonymous SNPs within individual genes. Linear regression of LD_nonsyn_ on LD_nsyn_ is shown as the red line. To control for the gene length, only SNPs within 300 nucleotides from each other were analyzed. Genes with fewer than 100 such pairs of SNPs were excluded. (**d**,**e**) The positive correlation between p_n_/p_s_ of the gene and its average LD (**d**) or the difference between LD_nonsyn_ and LD_syn_ (**e**). Here, the data on the USA population of *S. commune* are shown; similar patterns in the Russian population are shown in Supplementary Fig. 10.

There is no one-to-one correspondence between haploblocks and genes, which are, on average, longer. Still, different genes are covered by haploblocks to different extent, which leads to wide variation in the strength of LD and other characteristics among them. The excess of LD_nonsyn_ over LD_syn_ is also largely restricted to the genes with high LD, *i*.*e*. those that contain haploblocks (Fig. 3c). As a result, because both haplotypes tend to be common in a haploblock (Fig. 3), this excess is much stronger for loci with MAF > 0.05.

LD between alleles of all kinds is higher within genes with large ratios of nonsynonymous and synonymous polymorphisms p_n_/p_s_ (Spearman correlation p-value < 2e-16, Fig. 3d). Genes with elevated p_n_/p_s_ also have a stronger excess of LD_nonsyn_ over LD_syn_ (Fig. 3e, Spearman correlation p-value = 4.4e-17). This excess is the strongest for genes with high overall LD, but its correlation with p_n_/p_s_ holds even when the overall LD is controlled for (Supplementary Fig. 11).

## Excess of LD_nonsyn_ requires stable polymorphism

Simulations show that positive epistasis alone cannot lead to the observed large excess LD_nonsyn_ over LD_syn_, for which two extra conditions need to be satisfied. The general reason for this is simple: in order for a substantial LD between not-too-rare alleles to appear, these alleles must persist in the population for a long enough time.

First, positive epistasis must lead to a full compensation of deleterious effects of individual alleles. In other words, the fitnesses of at least two most-fit genotypes that are present in the population at substantial frequencies must be (nearly) the same (Supplementary Fig. 12). If this is not the case, selection favoring the only most-fit genotype leads to a too low level of genetic variation, which persists only due to recurrent mutation. It is natural to assume that the two major haplotypes that are common within a haploblock correspond to high-fitness genotypes. High-fitness genotypes can represent either isolated fitness peaks of equal heights (corresponding to a situation when two out of the four allele combinations confer high fitness) or a flat, curved ridge of high fitness (corresponding to a situation when three out of four combinations confer high fitness). The available data are insufficient to distinguish between these two options. Of course, with complete selective neutrality of all allele combinations there is no reason for LD_nonsyn_ > LD_syn_, so that at least some mixed genotypes, carrying alleles from different high-fitness genotypes, must be maladapted.

Second, there must be some kind of balancing selection that specifically works to maintain variation, because otherwise random drift does not allow genetic variation to persist for a long enough time even if some, or even all, genotypes are equally fit (Supplementary Fig. 12). Here, there are at least two options. On the one hand, a *bona fide* negative frequency-dependent selection (NFDS) can act either directly at loci that display high LD or at some other tightly linked loci^40,41^. On the other hand, variation can be maintained due to associative overdominance (AOD), resulting from selection against recurrent deleterious mutations at linked loci^42–44^.

Balancing selection is also a *sine qua non* for the presence of haploblocks, because a pair of divergent haplotypes can evolve in a panmictic population only if they coexist for a considerable time. A single locus under NFDS is enough to maintain a haploblock comprising the region of the genome around it. By contrast, if variation is maintained by AOD, it is more likely that selection against recessive mutations acts at a number of tightly linked loci^44^. Long coexistence of diverged haplotypes that comprise a haploblock enables accumulation of co-adapted combinations of nonsynonymous alleles within them. Thus, it is not surprising that a pronounced excess of LD_nonsyn_ over LD_syn_ in *S. commune* is observed primarily within haploblocks and that this excess is higher in genes with higher p_n_/p_s_.

## Correlated LDs in two populations

Although a high excess of LD_nonsyn_ is observed only within haploblocks, a signature of epistasis can also be seen outside of them in the form of a correlation between LDs in the two populations. This correlation can be high even if LDs *per se* are low.

The USA and the Russian populations share a large proportion of their SNPs. Given the high divergence between the two populations, few such shared SNPs are expected to have common origin in the ancestral population, and instead they are likely to have arisen from recurrent mutation. The high prevalence of coincident SNPs is not surprising because SNPs comprise 0.28 and 0.13 of all the aligned nucleotide sites in the USA and Russian populations, respectively^3^, Supplementary Fig. 2). We identified pairs of shared biallelic SNPs located within 2kb from one another and calculated the LD between them in both populations. To avoid the effects of strong within-population linkage and the occasional co-occurrence of haploblocks between populations, we excluded SNPs located within haploblocks or within genes under high LD (> 0.8 LD quantile for the corresponding population) in either population.

Values of LD in the two populations are strongly correlated only for pairs of nonsynonymous SNPs located within the same gene, and only if both populations carry the same pairs of amino acids in the same sites (Fig. 4). Correlation of LDs is the strongest if shared SNPs carry the same pairs of nucleotides, but is also observed if they encode the same amino acids by different nucleotides (Supplementary Fig. 13). The contrast between correlations within pairs of sites that reside in the same vs. different genes and the correlation of LDs observed for different nucleotides encoding the same amino acid cannot be explained by inheritance of LD from the common ancestral population. Moreover, synonymous SNPs are expected to be on average older than nonsynonymous ones, so that this mechanism should lead to a higher correlation of LDs for pairs of synonymous sites. Thus, the observed pattern indicates that epistatic selection is shared between the two populations.

**Figure 4.**
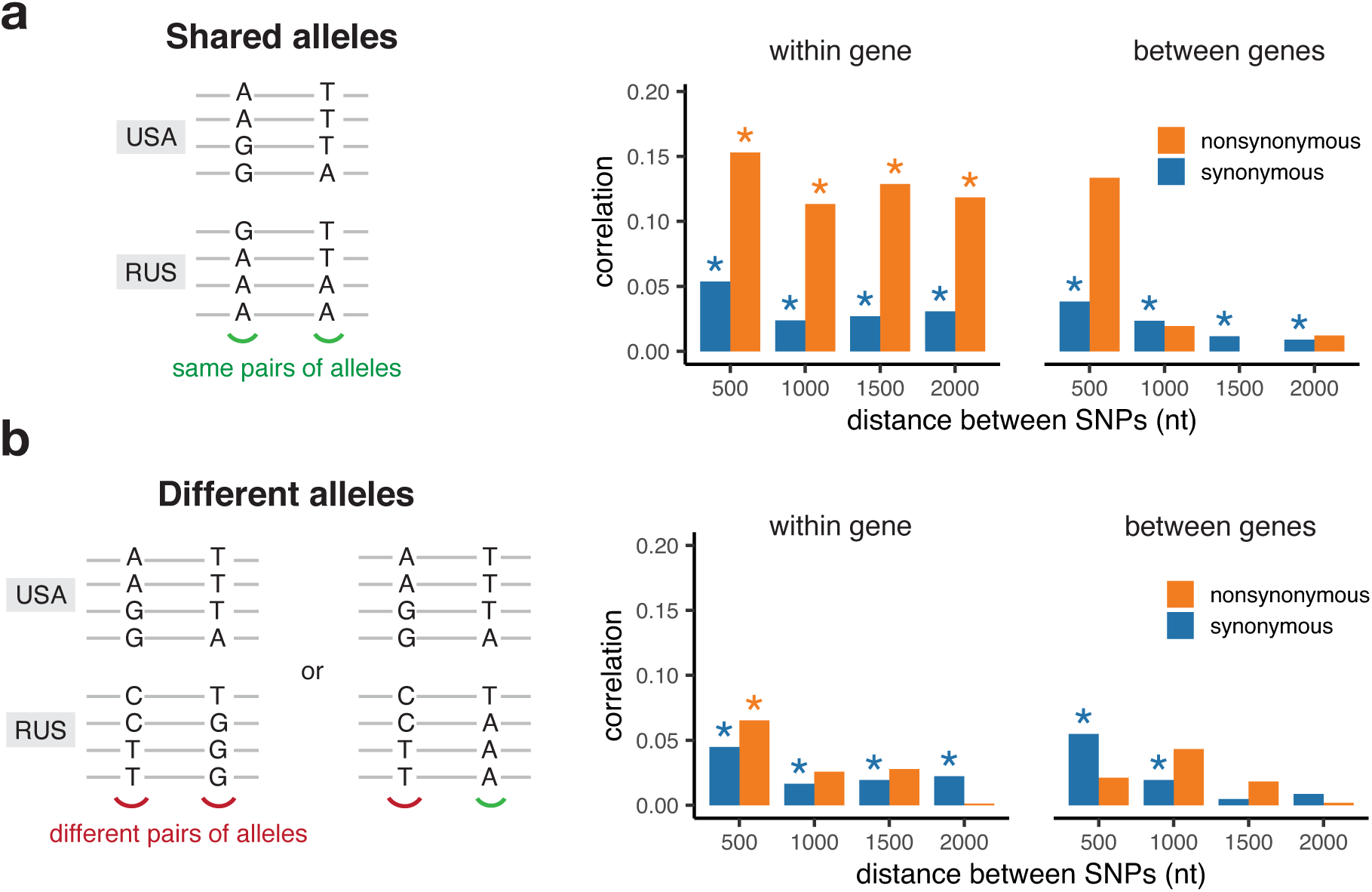
Correlation of LD values between pairs of shared SNPs in the two *S. commune* populations. (**a**) Pairs of SNPs with the same alleles in both sites, (**b**) pairs of SNPs differing by at least one allele. Asterisks indicate Spearman correlation p-values < 0.001.

Correlation of LDs between SNPs located within haploblocks in both populations is high regardless of whether they reside in the same or different genes, apparently because of occasional coincidence of haploblocks between populations (Supplementary Fig. 14).

## Discussion

On top of its most salient property, an exceptionally high π, genetic variation within *S. commune* possesses two other pervasive features. The first is a high prevalence of mostly short haploblocks, genome segments comprising two or occasionally three distinct haplotypes, which is a signature of balancing selection. The overall fraction of the genome covered by haploblocks is ∼10%, which is about an order of magnitude higher than the fraction covered by detectable signatures of balancing selection in genomes of other species^45–47^.

The second feature is the excessive attraction between nonsynonymous alleles polarized by frequency. This pattern is much stronger within haploblocks, indicating that they were shaped by both balancing and epistatic selection, so that amino acids common within a haplotype together confer a higher fitness. Polymorphisms that involve haplotypes that comprise many interacting genes, such as inversions^48–51^ and supergenes^52,53^, are known from the dawn of population genetics, but here we are dealing with an analogous phenomenon at a much finer scale, because haploblocks are typically shorter than genes. Thus, instead of coadapted gene complexes^48^, haplotypes represent coadaptive site complexes within genes.

In our simulations, equally high fitnesses of two or more genotypes was a necessary condition for a large excess of LD_nonsyn_, because otherwise the polymorphism did not live long enough for any substantial LD to evolve. However, epistasis between loci responsible for real or apparent balancing selection and those involved in compensatory interactions probably abolished the need for this fine-tuning of fitnesses. For example, if each haploblock carries its own complement of partially recessive deleterious mutations, together with alleles engaged in compensatory interactions with each other which also make these recessive mutations less deleterious, AOD can be expected to cause stable coexistence of these alleles.

Why are haploblocks and positive LD between rare nonsynonymous alleles so common in *S. commune*, but not in other, less polymorphic, species? There may be several, not mutually exclusive, reasons for this. Regarding haploblocks, real or apparent balancing selection may be more common in *S. commune* due to its higher polymorphism. Also, the same balancing selection may protect polymorphism in a huge population of *S. commune*, but not in populations with lower N_e_. Finally, an excess of haploblocks in *S. commune* may be at least due to better detection of signatures of balancing selection in a species with an extraordinary density of SNPs.

Excessive LD_nonsyn_ in *S. commune* is also likely to be due to its hyperpolymorphism which increases the probability that mutually compensating alleles at a pair of interacting sites achieve high frequency and encounter each other in the same haplotype before being eliminated by selection. In other words, even if the fitness landscape remains the same, it results in more epistatic selection and, thus, in stronger LD in a species whose genetic variation covers a larger chunk of this landscape (Fig. 1).

In a vast majority of species, π is a small parameter << 1. This imposes a severe constraint on operation of selection and obscures signatures of its particular modes. Thus, hyperpolymorphic species where π is ∼1 provide a unique opportunity to study phenomena which are traditionally viewed as belonging to the domain of macroevolution through data on within-population variation.

## Materials and methods

### *S. commune* sampling, sequencing and assembly

Haploid cultures of 24 isolates, each originated from a single haplospore, were obtained from fruit bodies collected in Ann Arbor, MI, USA by T. James and A. Kondrashov and in Moscow and Kostroma regions, Russia by A. Kondrashov, A. Baykalova and T. Neretina in 2009–2015. Specimen vouchers are stored in the White Sea Branch of Zoological Museum of Moscow State University (WS). Herbarium numbers are listed in Supplementary Table 2. To obtain isolates, wild fruit bodies were hung on the top lid of a 10 cm petri dish with agar medium. Petri dish was set at an angle of 60-70 degrees to the horizontal surface for 32 hours. A germinated spore was excised together with a square-shaped fragment (approximately 0.7×0.7 mm) of the medium from the maximally rarefied area of the obtained spore print under a stereomicroscope with 100x magnification. The obtained isolates were cultured in Petri dishes on 2% malt extract agar for a week. For storage, cultures were subcultured into 1.5 ml microcentrifuge tubes with 2% malt extract agar. To obtain sufficient biomass for DNA isolation, isolates were cultured in 20 ml 0.5% malt extract liquid medium in 50 ml microcentrifuge tubes in a horizontal position on a shaker at 100 rpm in daylight for 5 to 10 days. The tubes with the cultures were then centrifuged at 4000 rpm, and the supernatant was decanted. The resulting mycelium was lyophilized. DNA was extracted using Diamond DNA kit according to the manufacturer’s recommendations.

DNA libraries were constructed using the NEBNext Ultra II DNA Library Prep Kit kit by New England Biolabs (NEB) and the NEBNext Multiplex Oligos for Illumina (Index Primers Set 1) by NEB following the manufacturer’s protocol. The samples were amplified using 10 cycles of PCR. The constructed libraries were sequenced on Illumina NextSeq500 with paired-end read length of 151. The genomes were assembled *de novo* using SPAdes (v3.6.0)^54^; possible contaminations were removed using *blobology*^*55*^. Average N50 was ∼165kb for USA samples and ∼70kb for Russian samples (assembly statistics are provided in Supplementary Table 2).

Together with the 30 samples sequenced previously^3,56^, the obtained haploid genomes were aligned with TBA and *multiz*^*57*^ and projected onto the reference scaffolds^58^. Ortholog sequences were extracted on the basis of the reference genome annotation^58^ and realigned using *macse* codon-based aligner^59^. The alignments are available at https://makarich.fbb.msu.ru/astolyarova/schizophyllum_data/. Only the gap-free columns of the whole-genome alignment and the orthologs that were found in all 55 genomes were used for analysis. The total number of detected SNPs was 5.8 million for the USA population (82% of them biallelic, Supplementary Table 3) and 2.7 million for the Russian population (93% biallelic). 25% of the USA SNPs were shared with the Russian population (11% with the same major and minor alleles), and 53% of the Russian SNPs were shared with the USA population (23% with the same major and minor alleles, Supplementary Fig. 1).

The phylogeny of the sequenced genomes was reconstructed with RAxML^60^ (Supplementary Fig. 2). Nucleotide diversity (π) was estimated as the average frequency of pairwise nucleotide differences; π for different classes of sites is shown in Supplementary Fig. 1. Two samples from Florida (USA population) were excluded from the further analysis to minimize the possible effect of population structure.

Genome sequence data are deposited at DDBJ/ENA/GenBank under accession numbers JAGVRL000000000-JAGVSI000000000, BioProject PRJNA720428. Sequencing data are deposited at SRA with accession numbers SRR14467839-SRR14467862.

### Data on *H. sapiens* and *D. melanogaster* populations

We used polymorphism data from 1296 phased human genomes from African and European super-populations sequenced as part of the 1000 Genomes project^61^ (Supplementary Table 4). If several individuals from the same family were sequenced, we included only one of them. As a *D. melanogaster* dataset, we used 197 haploid genomes from the Zambia population^62^. Only autosomes were analyzed in both datasets.

### Estimation of LD

As a measure of linkage disequilibrium between two biallelic sites, we used r^2^, calculated as follows:

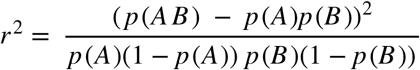, where p(A) and p(B) are the minor allele frequencies at these sites, and p(AB) is the frequency of the genotype that carries minor alleles at both sites.

Singletons (sites with minor allele present only in one genotype) were excluded from the analysis.

### Haploblocks annotation

In order to annotate the haploblocks, we calculated LD along the *S. commune* genome in a sliding window of 250 nucleotides with a step of 20 nucleotides (only non-singleton SNPs are analyzed; the windows with less than 10 SNPs were excluded). Any continuous sequence of overlapping windows with r^2^ larger than the threshold value was merged together in a haploblock. The LD threshold value was defined independently for each *S. commune* population as the heavy tail of the within-window LD distribution, as compared with the lognormal distribution with the same mean and variance as in the data (Supplementary Fig. 15).

### Estimation of LD between physically interacting amino acid sites

Of 16,319 annotated protein-coding genes of *S. commune*^*58*^ 9,941 were found in all 55 aligned genomes. We blasted the protein sequences of these orthologous groups against the PDB database of protein structures. About 52% of them (5,188) had a match (e-value threshold = 1e-5) amongst the proteins with the known structure. We realigned the sequences of *S. commune* protein and the matching PDB protein with *clustal* and calculated within-population LD and physical distance (Å) for each pair of aligned positions. A pair of amino acid sites was considered physically adjacent if they were located within 10 Å from each other.

To compare LD between pairs of physically close and distant sites, we used the controlled permutation test (Fig. 2a): for each pair of physically close amino acid sites (within 10Å) we sampled a pair of physically distant amino acids on the same genetic distance (measured in aa). Pairs of sites closer than 5 aa were excluded from the analysis. To examine LD patterns within individual protein structures, we calculated contingency tables of pairs of SNPs being located in codons encoding physically close amino acids and having high LD (no less than 90% quantile for a given gene). Pairs of amino acid sites located closer than 30 aa or more distant than 100 aa from each other were excluded; genes with less than 5 pairs of physically close sites under high or low LD were also excluded. From these contingency tables, we calculated the odds ratio (OR) and chi-square test p-value for each gene. p-values were adjusted using BH correction. We identified 22 genes with pairs of adjacent sites having significantly higher LD in the USA population (out of 1,286 eligible genes in total), and 87 genes in the Russian population (out of 967) under 5% FDR (Supplementary Table 1). Examples of such genes are shown in Fig. 2 and Supplementary Fig. 5-6.

### Simulations of epistasis

To simulate evolution of populations with or without epistasis and balancing selection (Supplementary Fig. 12), we used an individual-based model implemented by *SLiM*^*63*^. Simulations were performed with diploid population size N=1000 and recombination rate 0. To achieve the level of genetic diversity π similar to *S. commune*, mutation rate µ was scaled as µ=π/2N=5e-5). The length of the simulated sequence was 100 nt. Each simulation started with a monomorphic population and proceeds for 100N generations. For calculations of synonymous and nonsynonymous LD, random 100 haploid genotypes were sampled from the population. Only SNPs with minor allele frequency > 5% in the sample were analyzed.

We modelled two types of sites, depending on whether mutations in them were neutral (with selection coefficient s_syn_ = 0) or weakly deleterious (s_nonsyn_ ≤ 0), representing synonymous and nonsynonymous sites correspondingly. There were twice as many nonsynonymous as synonymous sites. Under the non-epistatic model, *s* was independent of the genetic background. We assumed s_nonsyn_=-0.01 with the dominance coefficient *h* of 0.5.

Under the pairwise positive epistasis model, we assumed that a mutation at one nonsynonymous site can be partially or fully compensated by a mutation at another site. In this model, all nonsynonymous sites were split into pairs. Each mutation of a pair individually occurring within a genotype was assumed to be deleterious, with selection coefficient s_nonsyn_=-0.01; however, the fitness of the double mutant is larger than expected under the additive (non-epistatic) model. We used several models of epistasis, with different strengths of epistasis strength and landscape shapes (Supplementary Fig. 12).

In the NFDS model of balancing selection, a single mutation at a random position was subjected to frequency-dependent selection (so that it is positively selected at frequencies below 0.5, and negatively selected at frequencies above 0.5). In the AOD model, mutations in 10 random positions were fully recessive (*h*=0) and weakly deleterious (*s*=-0.0025).

To simulate evolution of populations with different levels of genetic diversity under epistasis (Supplementary Note 1), we used *FFPopSim*^*64*^. To achieve different levels of genetic diversity π, mutation rate µ was scaled as µ=π/2N. The calculations were performed the same way as in *SLiM*, but In this case, we used haploid population size N=2000, population-scaled recombination rate 0.01 and the simulated sequence length of 300 nucleotides.

## Acknowledgements

We thank Timothy James, Anna Baykalova and members of Bazykin and Kondrashov labs for collecting *S. commune* samples. We thank Shamil Sunyaev and members of his lab for useful comments on drafts of this article.

## Author contributions

G.A.B. conceived the study; A.S.K., T.V.N. and G.A.B. performed sample collection; E.A.Z. and T.V.N. cultivated *S. commune* and extracted DNA; A.V.F. constructed Illumina libraries and performed the Illumina sequencing; A.V.S., G.A.B. and A.S.K. designed the analyses; A.V.S. analyzed the data and performed simulations; A.S.K., G.A.B., and A.V.S. wrote the paper with contributions from all authors.

## References

1. Leffler, E. M. et al. Revisiting an old riddle: what determines genetic diversity levels within species? PLoS Biol. 10, e1001388 (2012).

2. Dey, A., Chan, C. K. W., Thomas, C. G. & Cutter, A. D. Molecular hyperdiversity defines populations of the nematode Caenorhabditis brenneri. Proc. Natl. Acad. Sci. U. S. A. 110, 11056–11060 (2013).

3. Baranova, M. A. et al. Extraordinary Genetic Diversity in a Wood Decay Mushroom. Mol. Biol. Evol. 32, 2775–2783 (2015).

4. Smith, J. M. Natural selection and the concept of a protein space. Nature 225, 563–564 (1970).

5. Gillespie, J. H. The Causes of Molecular Evolution. (Oxford University Press, 1994).

6. Povolotskaya, I. S. & Kondrashov, F. A. Sequence space and the ongoing expansion of the protein universe. Nature 465, 922–926 (2010).

7. de Visser, J. A. G. M., Cooper, T. F. & Elena, S. F. The causes of epistasis. Proc. Royal Soc. B 278, 3617–3624 (2011).

8. McCandlish, D. M., Rajon, E., Shah, P., Ding, Y. & Plotkin, J. B. The role of epistasis in protein evolution. Nature vol. 497 E1–E2 (2013).

9. de Visser, J. A. G. M. & Krug, J. Empirical fitness landscapes and the predictability of evolution. Nat. Rev. Genet. 15, 480–490 (2014).

10. Good, B. H. & Desai, M. M. The impact of macroscopic epistasis on long-term evolutionary dynamics. Genetics 199, 177–190 (2015).

11. Kryazhimskiy, S., Dushoff, J., Bazykin, G. A. & Plotkin, J. B. Prevalence of epistasis in the evolution of influenza A surface proteins. PLoS Genet. 7, e1001301 (2011).

12. Dobzhansky, T. Studies on Hybrid Sterility. II. Localization of Sterility Factors in Drosophila Pseudoobscura Hybrids. Genetics 21, 113–135 (1936).

13. Orr, H. A. The population genetics of speciation: the evolution of hybrid incompatibilities. Genetics 139, 1805–1813 (1995).

14. Kondrashov, A. S., Sunyaev, S. & Kondrashov, F. A. Dobzhansky–Muller incompatibilities in protein evolution. Proc. Natl. Acad. Sci. U. S. A. 99, 14878–14883 (2002).

15. Sackton, T. B. & Hartl, D. L. Genotypic Context and Epistasis in Individuals and Populations. Cell 166, 279–287 (2016).

16. Crow, J. F. On epistasis: why it is unimportant in polygenic directional selection. Philos. Trans. R. Soc. Lond. B Biol. Sci. 365, 1241–1244 (2010).

17. Mäki-Tanila, A. & Hill, W. G. Influence of gene interaction on complex trait variation with multilocus models. Genetics 198, 355–367 (2014).

18. Hill, W. G., Goddard, M. E. & Visscher, P. M. Data and theory point to mainly additive genetic variance for complex traits. PLoS Genet. 4, e1000008 (2008).

19. Hivert, V., Sidorenko, J., Rohart, F., Goddard, M. E. & Yang, J. Estimation of non-additive genetic variance in human complex traits from a large sample of unrelated individuals. American journal of human genetics 108,5, 786–798 (2021).

20. Barton, N. H. Genetic linkage and natural selection. Philos. Trans. R. Soc. Lond. B Biol. Sci. 365, 2559–2569 (2010).

21. Takahasi, K. R. & Tajima, F. Evolution of coadaptation in a two-locus epistatic system. Evolution 59, 2324–2332 (2005).

22. Kouyos, R. D., Silander, O. K. & Bonhoeffer, S. Epistasis between deleterious mutations and the evolution of recombination. Trends Ecol. Evol. 22, 308–315 (2007).

23. Pedruzzi, G., Barlukova, A. & Rouzine, I. M. Evolutionary footprint of epistasis. PLoS Comput. Biol. 14, e1006426 (2018).

24. Boyrie, L., Moreau, C., Frugier, F., Jacquet, C. & Bonhomme, M. A linkage disequilibrium-based statistical test for Genome-Wide Epistatic Selection Scans in structured populations. Heredity 126, 77–91 (2021).

25. Wang, M.-C., Chen, F.-C., Chen, Y.-Z., Huang, Y.-T. & Chuang, T.-J. LDGIdb: a database of gene interactions inferred from long-range strong linkage disequilibrium between pairs of SNPs. BMC Res. Notes 5, 212 (2012).

26. Beissinger, T. M. et al. Using the variability of linkage disequilibrium between subpopulations to infer sweeps and epistatic selection in a diverse panel of chickens. Heredity 116, 158–166 (2016).

27. Zan, Y., Forsberg, S. K. G. & Carlborg, Ö. On the Relationship Between High-Order Linkage Disequilibrium and Epistasis. G3 8, 2817–2824 (2018).

28. Garcia, J. A. & Lohmueller, K. E. Negative linkage disequilibrium between amino acid changing variants reveals interference among deleterious mutations in the human genome. PLoS Genet. 17, 1–25 (2021).

29. Ochs, I. E. & Desai, M. M. The competition between simple and complex evolutionary trajectories in asexual populations. BMC Evol. Biol. 15, 55 (2015).

30. Dobzhansky, T. Genetics and the origin of species. 287 (Columbia University Press, 1937).

31. Bateson, W. Heredity and variation in modern lights. in Darwin and Modern Science (ed. Seward, A. C.) 85–101 (Cambridge University Press, 1909).

32. Muller, H. Isolating mechanisms, evolution, and temperature. Biol. Symp. 6, 71–125 (1942).

33. Gavrilets, S. Evolution and speciation on holey adaptive landscapes. Trends Ecol. Evol. 12, 307–312 (1997).

34. Wright, S. The roles of mutation, inbreeding, crossbreeding and selection in evolution. Proc 6th Int Cong Genet. 1, 356–366 (1932).

35. Crow, J. F. & Kimura, M. An introduction to population genetics theory. (New York, Evanston and London: Harper & Row, Publishers, 1970).

36. Seplyarskiy, V. B. et al. Crossing-over in a hypervariable species preferentially occurs in regions of high local similarity. Mol. Biol. Evol. 31, 3016–3025 (2014).

37. Ovchinnikov, S., Kamisetty, H. & Baker, D. Robust and accurate prediction of residue–residue interactions across protein interfaces using evolutionary information. Elife 3, e02030 (2014).

38. Marks, D. S. et al. Protein 3D structure computed from evolutionary sequence variation. PLoS One 6, e28766 (2011).

39. Sjodt, M. et al. Structure of the peptidoglycan polymerase RodA resolved by evolutionary coupling analysis. Nature 556, 118–121 (2018).

40. Charlesworth, D. Balancing selection and its effects on sequences in nearby genome regions. PLoS Genet. 2, e64 (2006).

41. Olendorf, R. et al. Frequency-dependent survival in natural guppy populations. Nature 441, 633–636 (2006).

42. Ohta, T. Associative overdominance caused by linked detrimental mutations. Genet. Res. 18, 277–286 (1971).

43. Zhao, L. & Charlesworth, B. Resolving the Conflict Between Associative Overdominance and Background Selection. Genetics 203, 1315–1334 (2016).

44. Gilbert, K. J., Pouyet, F., Excoffier, L. & Peischl, S. Transition from Background Selection to Associative Overdominance Promotes Diversity in Regions of Low Recombination. Curr. Biol. 30, 101–107.e3 (2020).

45. DeGiorgio, M., Lohmueller, K. E. & Nielsen, R. A model-based approach for identifying signatures of ancient balancing selection in genetic data. PLoS Genet. 10, e1004561 (2014).

46. Leffler, E. M. et al. Multiple instances of ancient balancing selection shared between humans and chimpanzees. Science 339, 1578–1582 (2013).

47. Rasmussen, M. D., Hubisz, M. J., Gronau, I. & Siepel, A. Genome-wide inference of ancestral recombination graphs. PLoS Genet. 10, e1004342 (2014).

48. Dobzhansky, T. & Pavlovsky, O. interracial hybridization and breakdown of coadapted gene complexes in Drosophila paulistorum and Drosophila willistoni. Proc. Natl. Acad. Sci. U. S. A. 44, 622–629 (1958).

49. Charlesworth, B. & Charlesworth, D. Selection of new inversions in multi-locus genetic systems. Genet. Res. 21, 167–183 (1973).

50. Singh, B. N. Chromosome inversions and linkage disequilibrium in Drosophila. Curr. Sci. 94, 459–464 (2008).

51. Sturtevant, A. H. & Mather, K. The Interrelations of Inversions, Heterosis and Recombination. Am. Nat. 72, 447–452 (1938).

52. Mather, K. The Genetical Architecture of Heterostyly in Primula sinensis. Evolution 4, 340–352 (1950).

53. Joron, M. et al. Chromosomal rearrangements maintain a polymorphic supergene controlling butterfly mimicry. Nature 477, 203–206 (2011).

54. Bankevich, A. et al. SPAdes: a new genome assembly algorithm and its applications to single-cell sequencing. J. Comput. Biol. 19, 455–477 (2012).

55. Kumar, S., Jones, M., Koutsovoulos, G., Clarke, M. & Blaxter, M. Blobology: exploring raw genome data for contaminants, symbionts and parasites using taxon-annotated GC-coverage plots. Front. Genet. 4, 237 (2013).

56. Bezmenova, A. V. et al. Rapid accumulation of mutations in growing mycelia of a hypervariable fungus Schizophyllum commune. Mol. Biol. Evol. (2020) doi:10.1093/molbev/msaa083.

57. Blanchette, M. et al. Aligning multiple genomic sequences with the threaded blockset aligner. Genome Res. 14, 708–715 (2004).

58. Ohm, R. A. et al. Genome sequence of the model mushroom Schizophyllum commune. Nat. Biotechnol. 28, 957–963 (2010).

59. Ranwez, V., Harispe, S., Delsuc, F. & Douzery, E. J. P. MACSE: Multiple Alignment of Coding SEquences accounting for frameshifts and stop codons. PLoS One 6, e22594 (2011).

60. Stamatakis, A. RAxML version 8: a tool for phylogenetic analysis and post-analysis of large phylogenies. Bioinformatics 30, 1312–1313 (2014).

61. 1000 Genomes Project Consortium et al. A global reference for human genetic variation. Nature 526, 68–74 (2015).

62. Lack, J. B. et al. The Drosophila genome nexus: a population genomic resource of 623 Drosophila melanogaster genomes, including 197 from a single ancestral range population. Genetics 199, 1229–1241 (2015).

63. Haller, B. C. & Messer, P. W. SLiM 3: Forward Genetic Simulations Beyond the Wright–Fisher Model. Mol. Biol. Evol. 36, 632–637 (2019).

64. Zanini, F. & Neher, R. A. FFPopSim: an efficient forward simulation package for the evolution of large populations. Bioinformatics 28, 3332–3333 (2012).

