## Supplementary Information for "Complex fitness landscape shapes variation in a hyperpolymorphic species"

#### Supplementary Notes

#### Supplementary Note 1
#### Epistatic selection is more efficient in genetically diverse populations

Genetic interactions affect operation of selection only in a sufficiently variable population. The potency of any kind of selection increases with the amount of variation; for epistatic selection, however, this increase is expected to be faster than linear, because it depends on the number of possible allele combinations. In a highly polymorphic population, a particular allele is more likely to co-occur in the same haplotype with an interacting, *e.g.*, compensatory, allele (Supplementary Note 1 Fig. 1a) which should increase the impact of epistasis on linkage disequilibrium.

To illustrate this point, we modelled the evolution of a genome region in the presence and in the absence of positive epistasis in a panmictic population. We assumed that all mutations at a set of sites are individually deleterious, and that all these sites are involved in pairwise positive (*i. e*., antagonistic) sign epistasis; specifically, each deleterious mutation can be fully compensated by another mutation at exactly one site elsewhere in the genome, which is also deleterious when present alone. In the non-epistatic simulations, the effects of mutations were independent; however, at the end of the simulation we randomly assigned the “interacting” pairs of sites to account for the random coincidence of deleterious alleles. We found that in this model a higher polymorphism increases the probability that a deleterious mutation is compensated before being eliminated by selection (Supplementary Note 1 Fig. 1b). This probability increases with genetic diversity even for the non-epistatic simulations, because increased diversity elevates the likelihood of randomly encountering a compensating allele in the same haplotype. For epistatic simulations, however, this increase is more radical, reflecting the effect of epistatic selection favoring compensated haplotypes.

After the mutation-selection equilibrium was reached, we measured the strength of epistatic selection between all segregating polymorphisms, asking to what extent the mutational load is reduced by epistasis maintaining combinations of compensatory mutations. As shown in Supplementary Note 1 Fig. 1c, the ability of epistatic selection to reduce the mutation load (*i. e*., to increase the mean fitness) strongly depends on $\pi$. In less variable populations ($\pi < 0.01$), epistasis is practically inefficient and doesn’t affect LD (Wilcoxon test p-values > 0.33); this is because the probability of occurrence of the favorable combination of alleles in the population for selection to act upon is low. In more diverse populations, however, such combinations may arise and be favored by epistatic selection, which increases LD between them (Wilcoxon test p-value < 0.01 for $\pi \geq 0.01$).

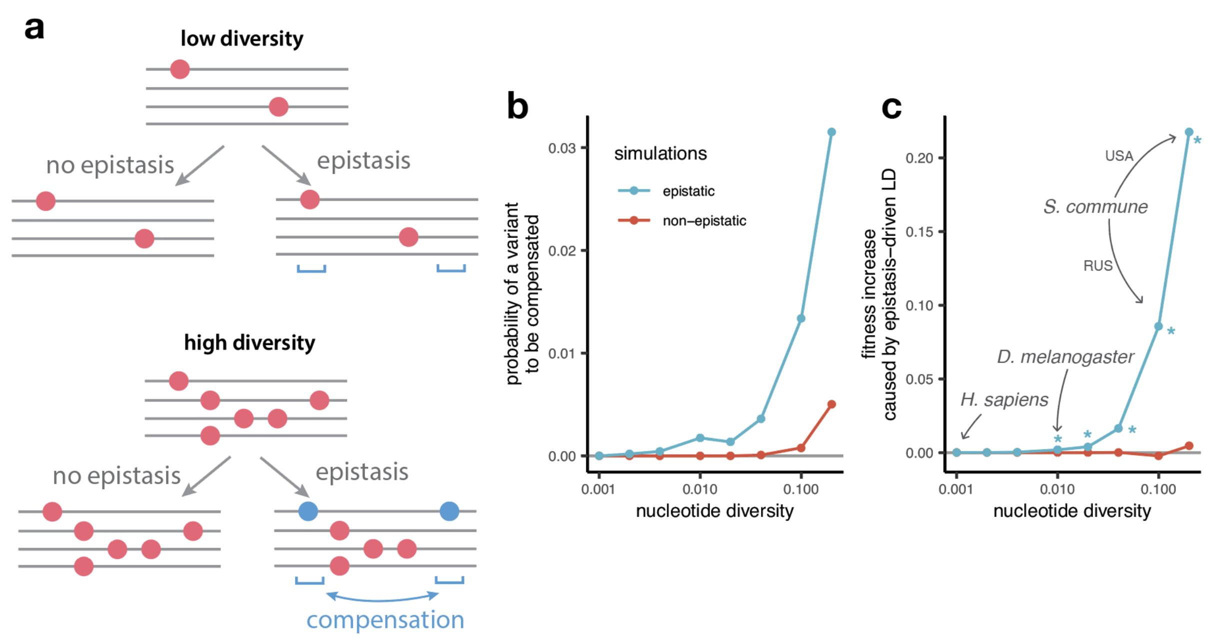


**Supplementary Note 1 Figure 1. The efficiency of epistasis in populations with different levels of nucleotide diversity.** (**a**) Under low nucleotide diversity, deleterious mutations (red dots) are unlikely to be compensated. If nucleotide diversity is high, epistatic selection maintains LD between SNPs in interacting sites (blue dots). (**b**) The probability that a deleterious variant is compensated by another variant within the same individual at the end of the simulation. (**c**) Increase in the mean fitness of the population caused by epistatic selection maintaining LD between favorable allele combinations. The fitness is plotted relative to that of a population consisting of individuals with uncorrelated alleles at different sites, obtained by permuting alleles among individuals. The efficiency of epistatic selection in maintaining linkage is much higher in genetically variable populations. Asterisks in (**c**) indicate significant deviation from 0 (Wilcoxon paired test p-value < 0.01). Each simulation was repeated between 100 and 10,000 times depending on genetic diversity.

### Supplementary Note 2
#### $LD_{nonsyn} > LD_{syn}$ requires positive epistasis

When alleles at different loci can be related to each other, it makes sense to consider the sign of both epistasis and LD. For example, if we are concerned with allele frequencies, all rare alleles can be viewed as analogous. Then, LD is positive (negative) if genotypes carrying both rare alleles are over-(under-)represented in the population, and epistasis is positive (negative) if such genotypes have fitnesses above (below) those expected if alleles act independently. Of course, overrepresentation (and excessive fitness) of combinations of rare alleles automatically entails the same for combinations of common alleles.

Although we report LD between pairs of polymorphic sites as $r^2$, which is symmetric regarding the major or minor variants, the observed high values of $r^2$ correspond to positive LD between minor alleles for both synonymous and nonsynonymous SNPs (Supplementary Note 2 Fig. 1-3). Thus, $LD_{nonsyn} > LD_{syn}$ means that attraction between minor nonsynonymous alleles is stronger than between minor synonymous alleles. This pattern may seem to be surprising, because there are three factors that work in the opposite direction.

First, random drift, which affects nearly-neutral synonymous sites more than nonsynonymous sites which are mostly under negative selection, leads to attraction between minor alleles[1]. Second, negative selection at nonsynonymous sites causes repulsion between rare, deleterious alleles, due to Hill-Robertson interference, even if this selection does not involve any epistasis[2–4]. Third, there are data on negative epistasis in this selection, which also should lead to repulsion of deleterious alleles and, thus, negative LD between rare nonsynonymous alleles[1,4,5]. The first and the second factors are weak and can produce noticeable LD only between tightly linked loci, while the third factor may generate even long-range LD. By contrast, $LD_{nonsyn} > LD_{syn}$ can be explained only by positive epistasis in selection at nonsynonymous sites.

Although negative selection generally results in $LD_{nonsyn} < LD_{syn}$, our simulations demonstrated that Hill-Robertson interference without epistasis can produce attraction between minor alleles under a rather restrictive set of conditions. In these simulations, weakly deleterious polymorphisms can achieve high frequency only in regions of low recombination, leading to $LD_{nonsyn} > LD_{syn}$ for extremely high MAF (Supplementary Note 2 Figure 4a). However, this effect doesn’t hold if assuming unequal fitness effects of deleterious mutations (Supplementary Note 2 Figure 4b) or while merging SNPs of different frequencies together (Supplementary Note 2 Figure 4c).

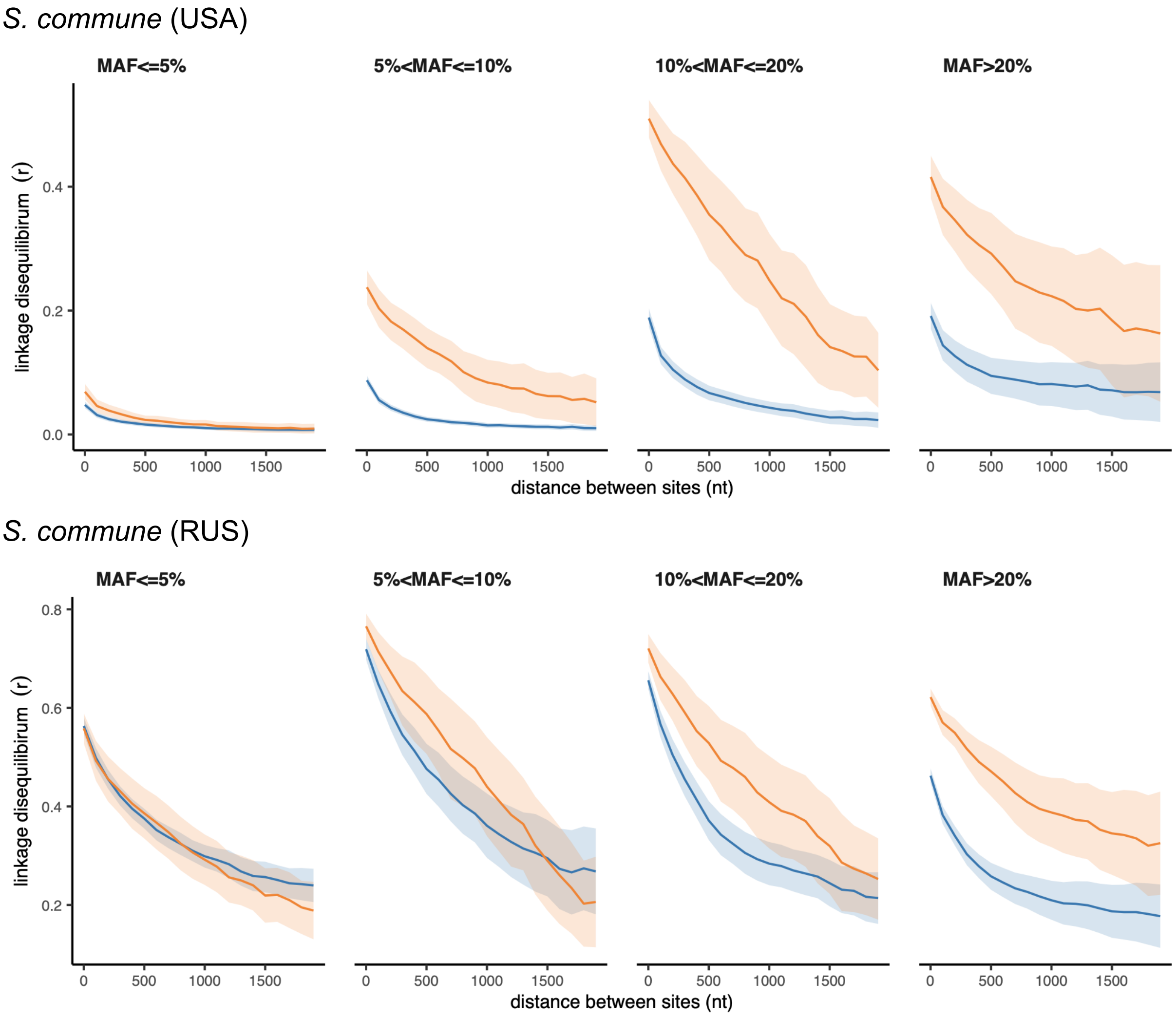


**Supplementary Note 2 Figure 1. Polarized linkage disequilibrium in *S. commune*.** LD between nonsynonymous SNPs is shown in orange, and LD between synonymous SNPs is shown in blue. Filled areas indicate SE of LD calculated for each scaffold separately.

*D. melanogaster*

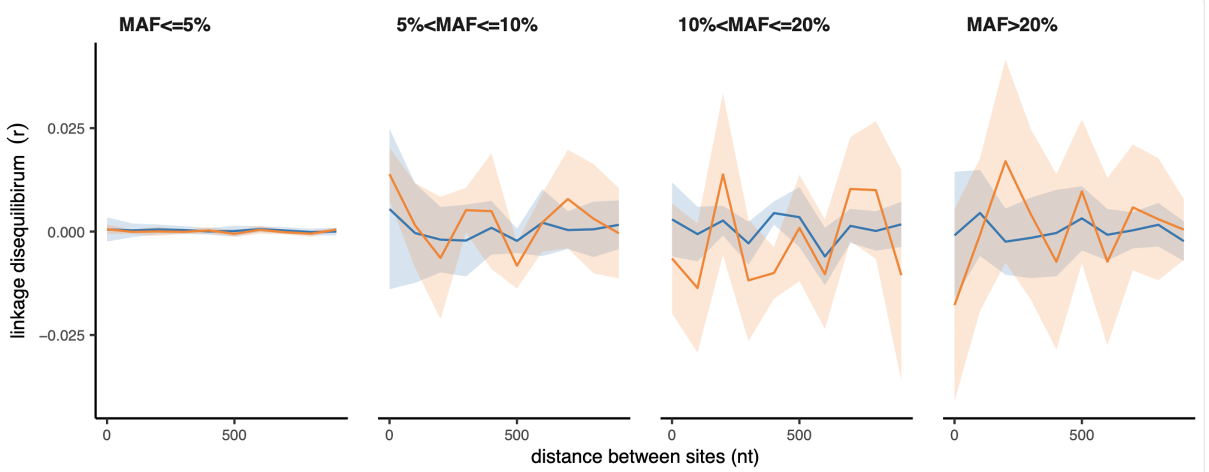


**Supplementary Note 2 Figure 2. Polarized linkage disequilibrium in *D. melanogaster*.** LD between nonsynonymous SNPs is shown in orange, and LD between synonymous SNPs is shown in blue. Filled areas indicate SE of LD calculated for each chromosome separately.

*H. sapiens*

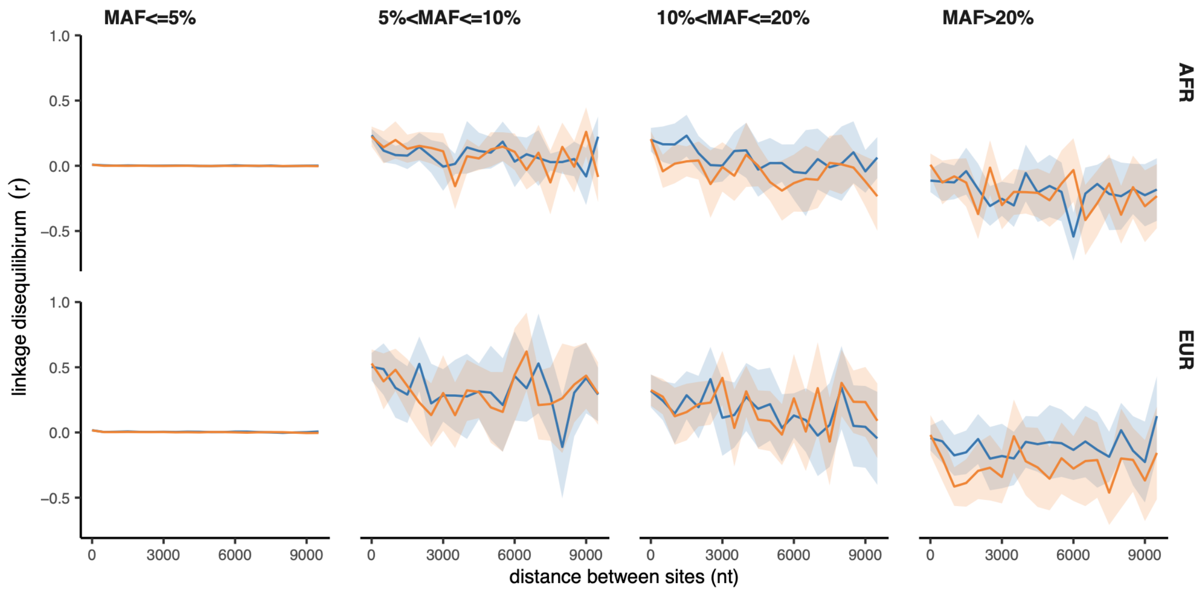


**Supplementary Note 2 Figure 3. Polarized linkage disequilibrium in *H.sapiens*.** LD between nonsynonymous SNPs is shown in orange, and LD between synonymous SNPs is shown in blue. Filled areas indicate SE of LD calculated for each chromosome separately.

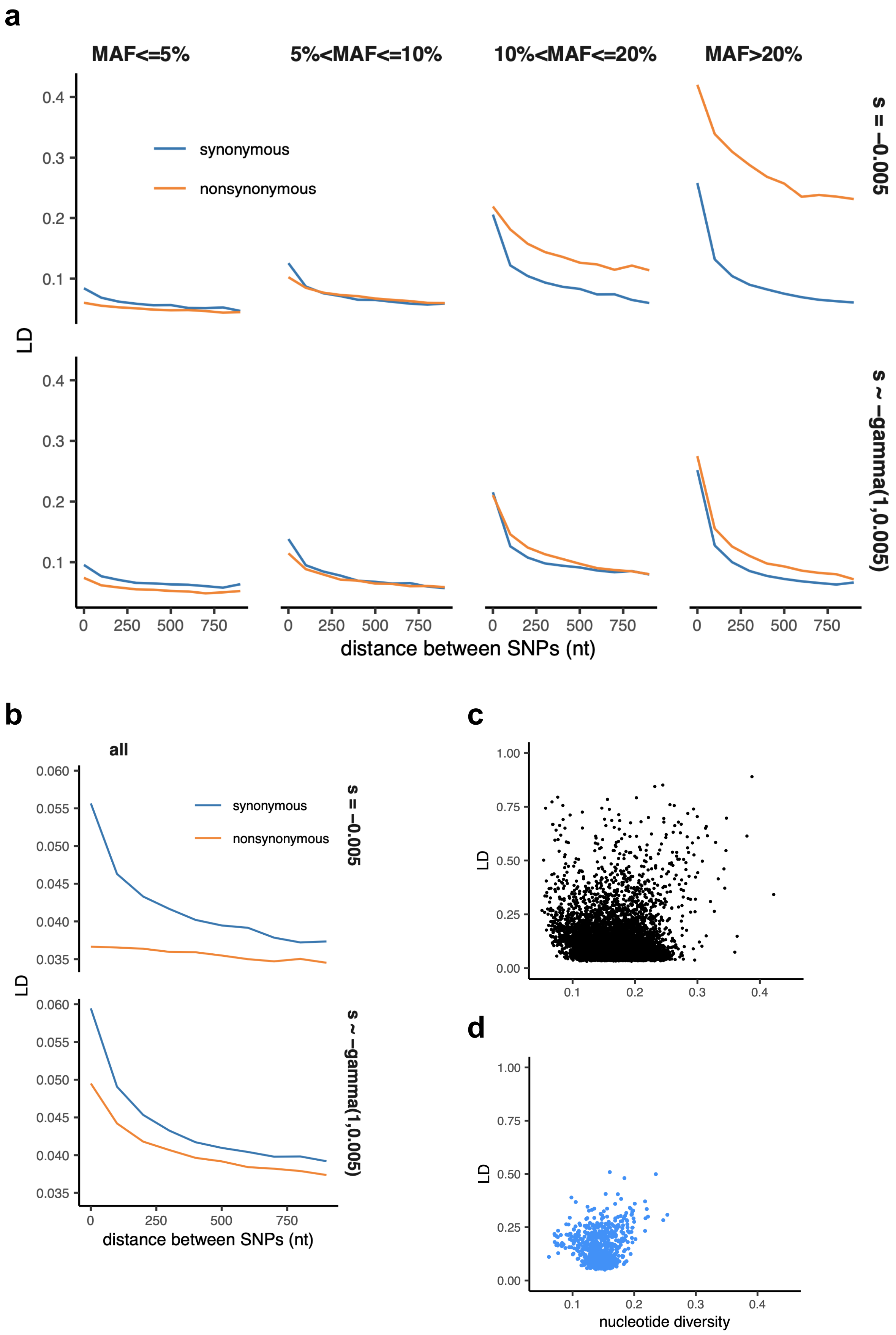
a
MAF<=5%
5%<MAF<=10%
10%<MAF<=20%
MAF>20%
synonymous
nonsynonymous
s = −0.005
s ~ −gamma(1,0.005)
LD
distance between SNPs (nt)
b
all
synonymous
nonsynonymous
s = −0.005
s ~ −gamma(1,0.005)
LD
distance between SNPs (nt)
c
LD
d
LD
nucleotide diversity

**Supplementary Note 2 Figure 4. Patterns of LD in simulations under negative selection.** (**a**) LD between nonsynonymous and synonymous pairs of SNPs split by MAF. (**b**) LD between all pairs of nonsynonymous and synonymous SNPs pooled together. (**a-b**) Haploid population size N = 2000, sequence length L = 1000 bp. Top panels - selection coefficients of all nonsynonymous mutations are equal to -0.005; bottom panels - selection coefficients of nonsynonymous mutations are gamma-distributed with parameters rate=1, scale=0.005. (**c**) LD and nucleotide diversity within genes of the USA population of *S. commune* (each point represents one gene). (**d**) LD and nucleotide diversity obtained in simulations.

#### Supplementary Note 3
#### Decreased $LD_{nonsyn}$ between low-frequency polymorphisms

The analysis of $LD_{nonsyn}$ and $LD_{syn}$ in the Main text (Fig. 1a-c) only considers high-frequency SNPs with MAF > 0.05. For *S. commune*, the excess $LD_{nonsyn}$ holds under different minor allele frequency thresholds (Supplementary Note 3 Fig. 1). However, in *D. melanogaster* and *H. sapiens,* rare nonsynonymous SNPs (with MAF < 0.05) taken alone show the opposite trend: the LD between such SNPs is reduced compared to synonymous SNPs at the same nucleotide distance (Supplementary Note 3 Fig. 2,3). In human populations, the vast majority of SNPs are rare, leading to $LD_{nonsyn} < LD_{syn}$ when all allele frequencies are considered (Supplementary Note 3 Fig. 3), in line with recently published results[4].

As discussed in Supplementary Note 2, decreased LD between negatively selected polymorphisms is expected due to Hill-Robertson interference between deleterious alleles[2,6]; this effect has been described previously for *H. sapiens*[4] and *D. melanogaster*[1] and is observed in our simulations (Supplementary Fig. 12). In addition, $LD_{nonsyn}$ can be reduced by negative epistasis between deleterious alleles[4], similarly to the negative LD detected among loss-of-function polymorphisms in humans, flies and plants[1,5].

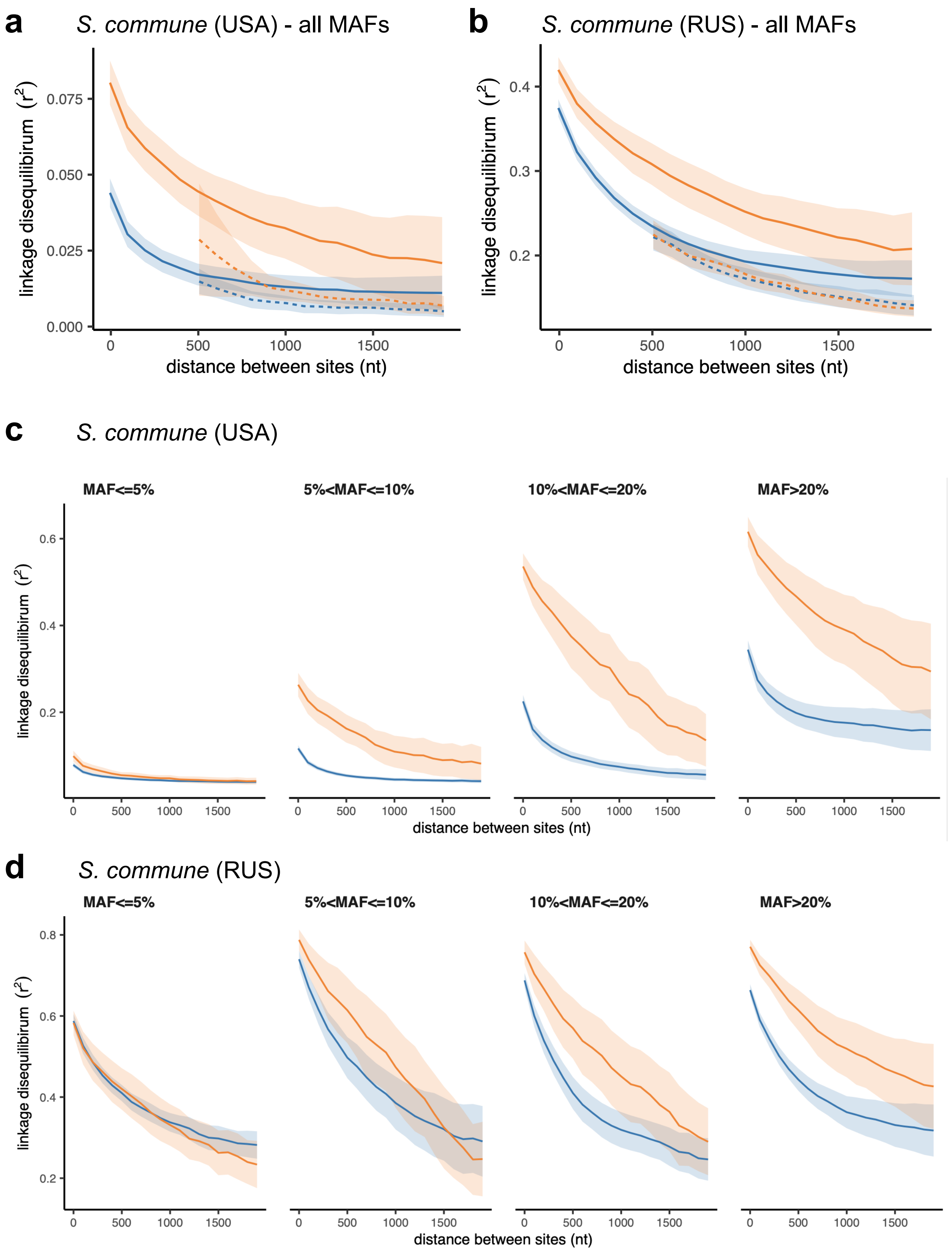


**Supplementary Note 3 Figure 1. LD between SNPs with different MAF in *S. commune*.** LD between nonsynonymous SNPs is shown in orange, and LD between synonymous SNPs is shown in blue. Filled areas indicate SE of LD calculated for each scaffold separately. (**a, b**) LD between all pairs of SNPs pooled together. Solid lines indicate LD between pairs of SNPs located within the same gene; dashed lines correspond to pairs of SNPs located in different genes. (**c, d**) Pairs of SNPs split by MAF.

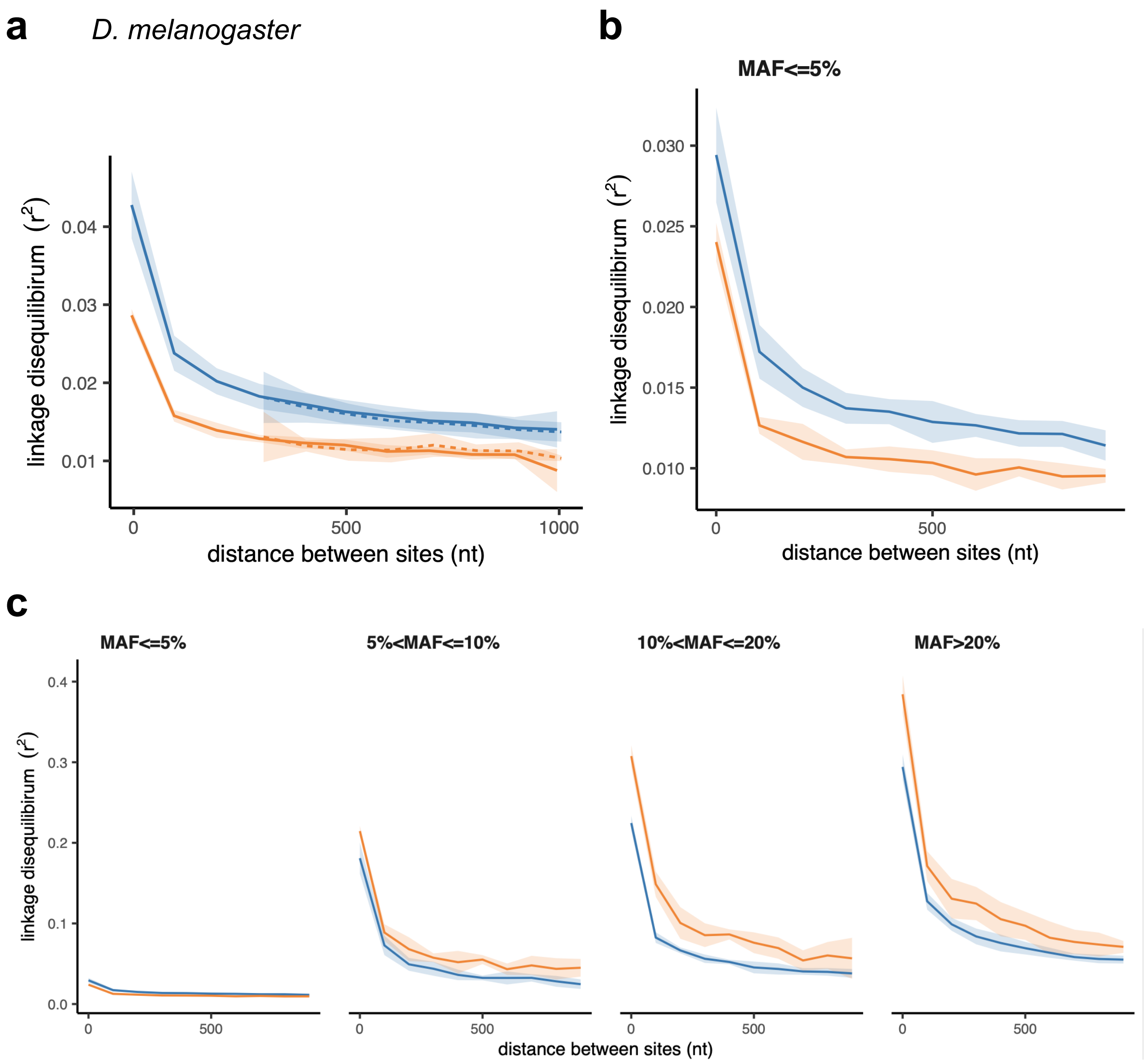


**Supplementary Note 3 Figure 2. LD between SNPs with different MAF in *D. melanogaster*.** LD between nonsynonymous SNPs is shown in orange, and LD between synonymous SNPs is shown in blue. Filled areas indicate SE of LD calculated for each chromosome separately. (**a**) LD between all pairs of SNPs pooled together. Solid lines indicate LD between pairs of SNPs located within the same gene; dashed lines correspond to pairs of SNPs located in different genes. (**b**) Pairs of SNPs with MAF < 0.05 (large scale). (**c**) Pairs of SNPs split by MAF.

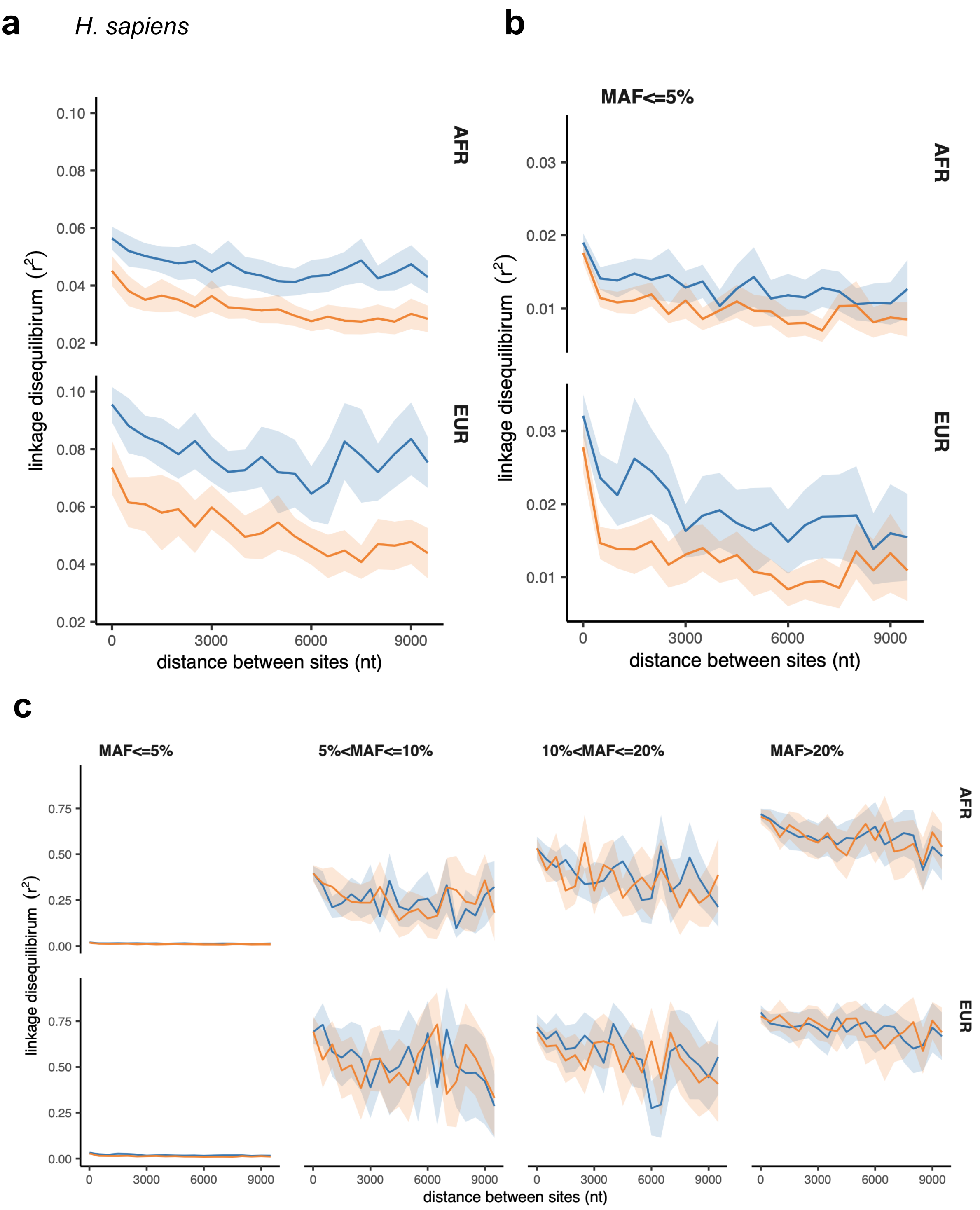


**Supplementary Note 3 Figure 3. LD between SNPs with different MAF in *H. sapiens*.** LD between nonsynonymous SNPs is shown in orange, and LD between synonymous SNPs is shown in blue. Filled areas indicate SE of LD calculated for each chromosome separately. (**a**) LD between all pairs of SNPs pooled together. Solid lines indicate LD between pairs of SNPs located within the same gene; dashed lines correspond to pairs of SNPs located in different genes. (**b**) Pairs of SNPs with MAF < 0.05 (large scale). (**c**) Pairs of SNPs split by MAF.

### References


1. Sandler, G., Wright, S. I. & Agrawal, A. F. Patterns and Causes of Signed Linkage Disequilibria in Flies and Plants. *Mol. Biol. Evol.* (2021).
2. Hill, W. G. & Robertson, A. The effect of linkage on limits to artificial selection. *Genet. Res.* **8**, 269–294 (1966).
3. Comeron, J. M., Williford, A. & Kliman, R. M. The Hill–Robertson effect: evolutionary consequences of weak selection and linkage in finite populations. *Heredity* **100**, 19–31 (2008).
4. Garcia, J. A. & Lohmueller, K. E. Negative linkage disequilibrium between amino acid changing variants reveals interference among deleterious mutations in the human genome. *PLoS Genet.* **17**, 1–25 (2021).
5. Sohail, M. *et al.* Negative selection in humans and fruit flies involves synergistic epistasis. *Science* **356**, 539–542 (2017).
6. Roze, D. & Barton, N. H. The Hill–Robertson Effect and the Evolution of Recombination. *Genetics* **173**, 1793–1811 (2006).

#### Supplementary Figures

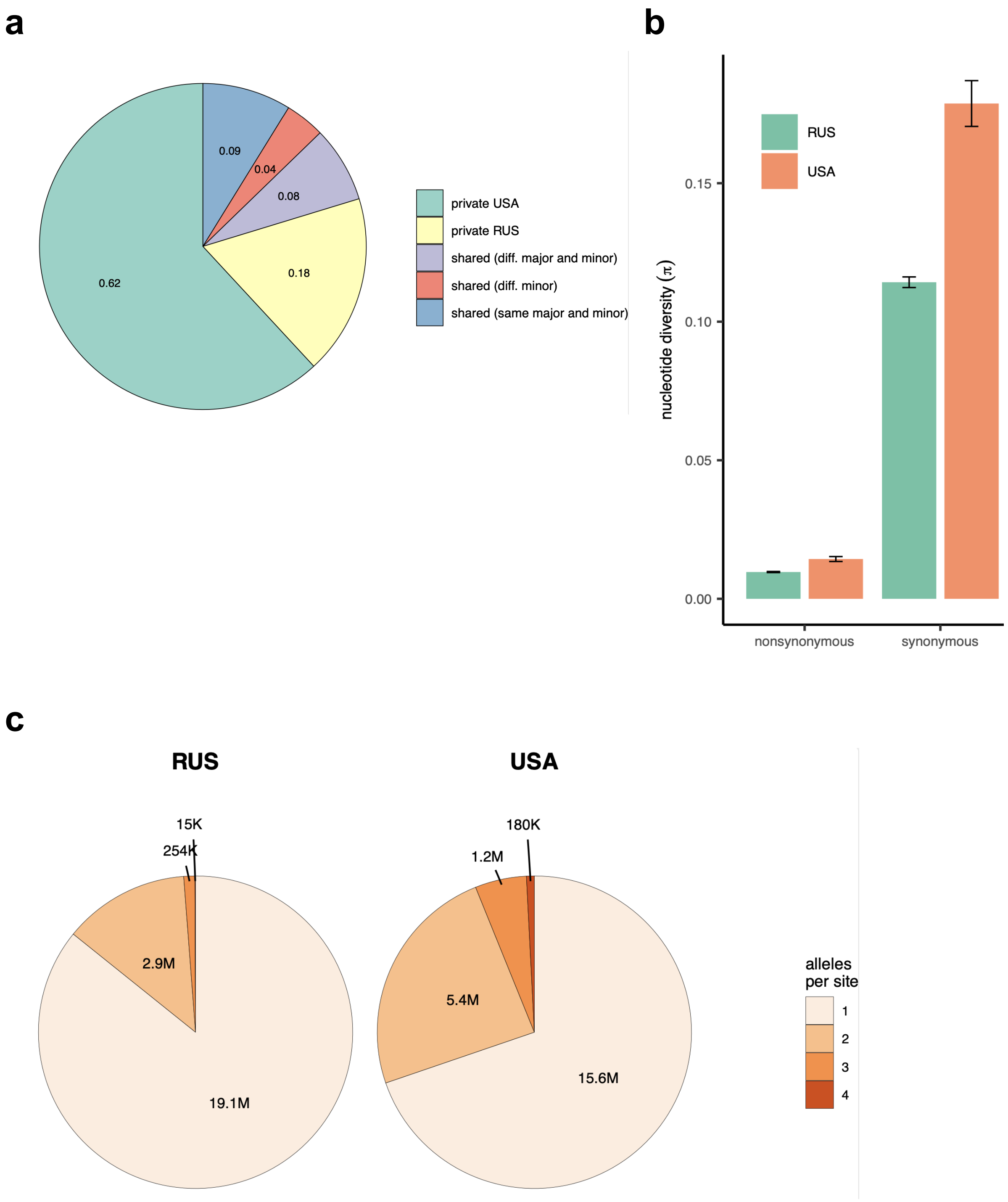


**Supplementary Figure 1. Patterns of nucleotide diversity in *S. commune*.** (**a**) The fraction of private and shared biallelic SNPs. (**b**) Within-population nucleotide diversity at different classes of sites (measured as π without Jukes-Cantor correction). (**c**) The number of monomorphic and polymorphic sites in the multiple whole-genome alignments of *S. commune* genomes.

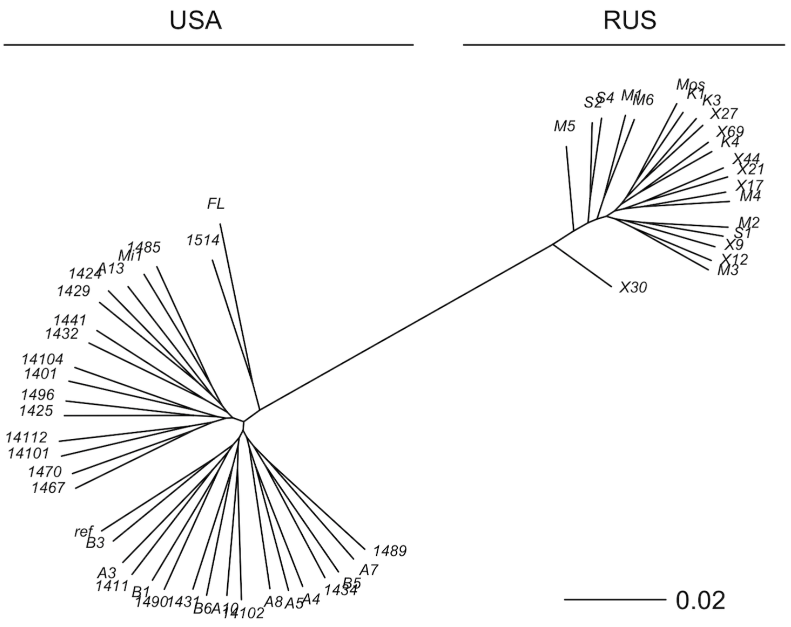


**Supplementary Figure 2. The reconstructed phylogeny of *S. commune*.**
USA and Russian populations of *S. commune* are highly divergent while having almost no within-population structure. Genetic distance is measured in nucleotide differences, the phylogeny is reconstructed based on the multiple whole-genome alignment.

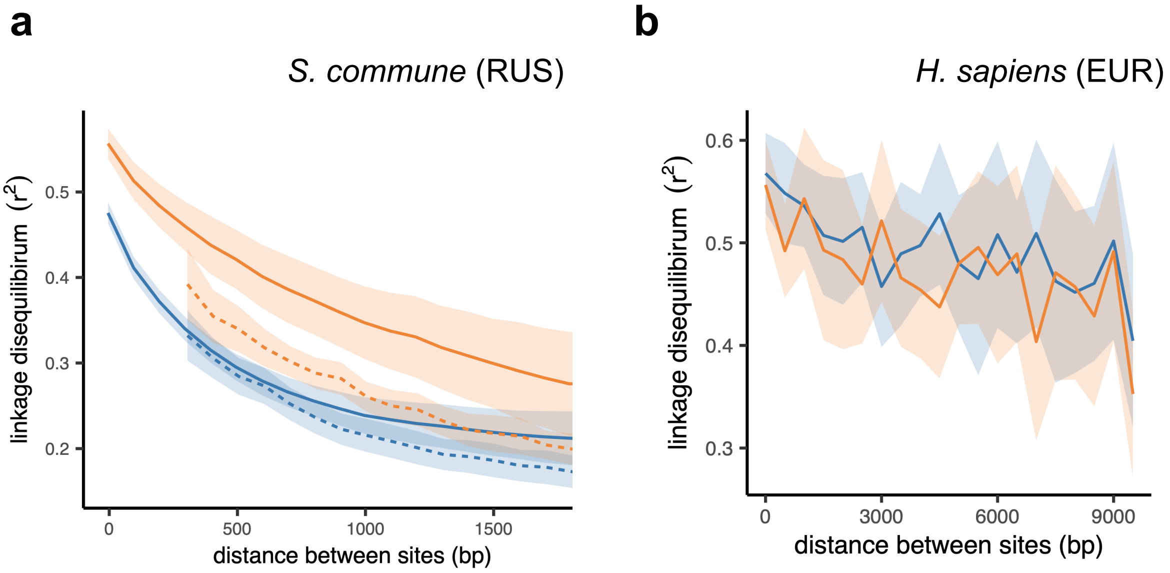


**Supplementary Figure 3. The efficiency of epistatic selection in populations with different levels of genetic diversity.** LD between nonsynonymous SNPs is shown in orange, and LD between synonymous SNPs is shown in blue. (**a**) Russian population of *S. commune*, (**b**) European super-population of *H. sapiens*. Solid lines indicate LD between pairs of SNPs located within the same gene; dashed lines correspond to pairs of SNPs located in different genes. Only SNPs with minor allele frequency > 0.05 are analysed. Filled areas indicate SE of LD calculated for each chromosome (for human) or scaffold (for *S. commune*) separately.

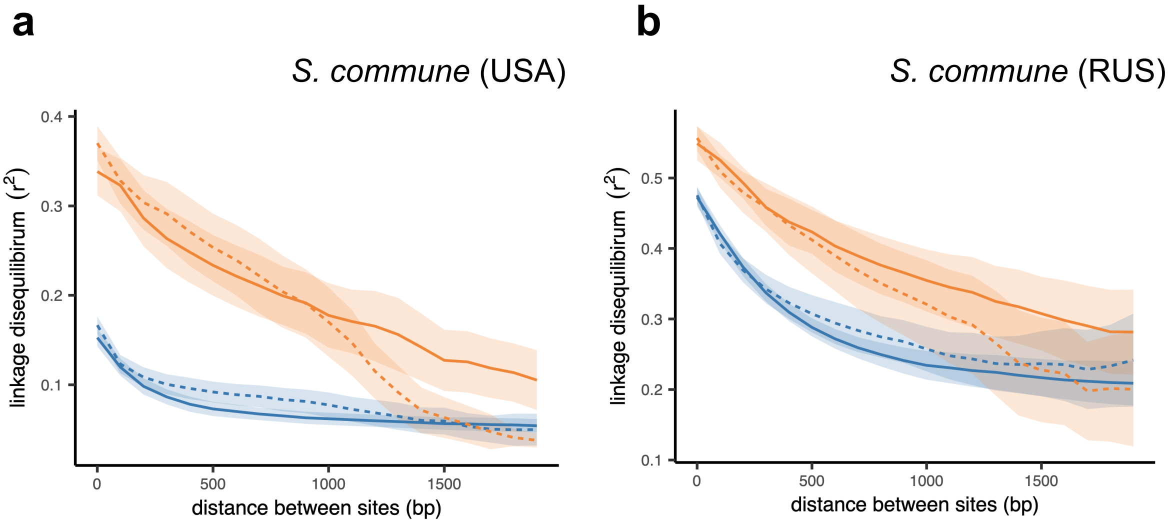


**Supplementary Figure 4. Linkage disequilibrium within and between exons in *S. commune*.** LD between nonsynonymous SNPs is shown in orange, and LD between synonymous SNPs is shown in blue. Solid lines indicate LD between pairs of SNPs located within the same exon of the gene; dashed lines correspond to pairs of SNPs located in different exons of the gene. (**a**) USA population of *S. commune*, (**b**) RUS population of *S. commune*. Only SNPs with minor allele frequency > 0.05 are analysed. Filled areas indicate SE of LD calculated for each scaffold separately.

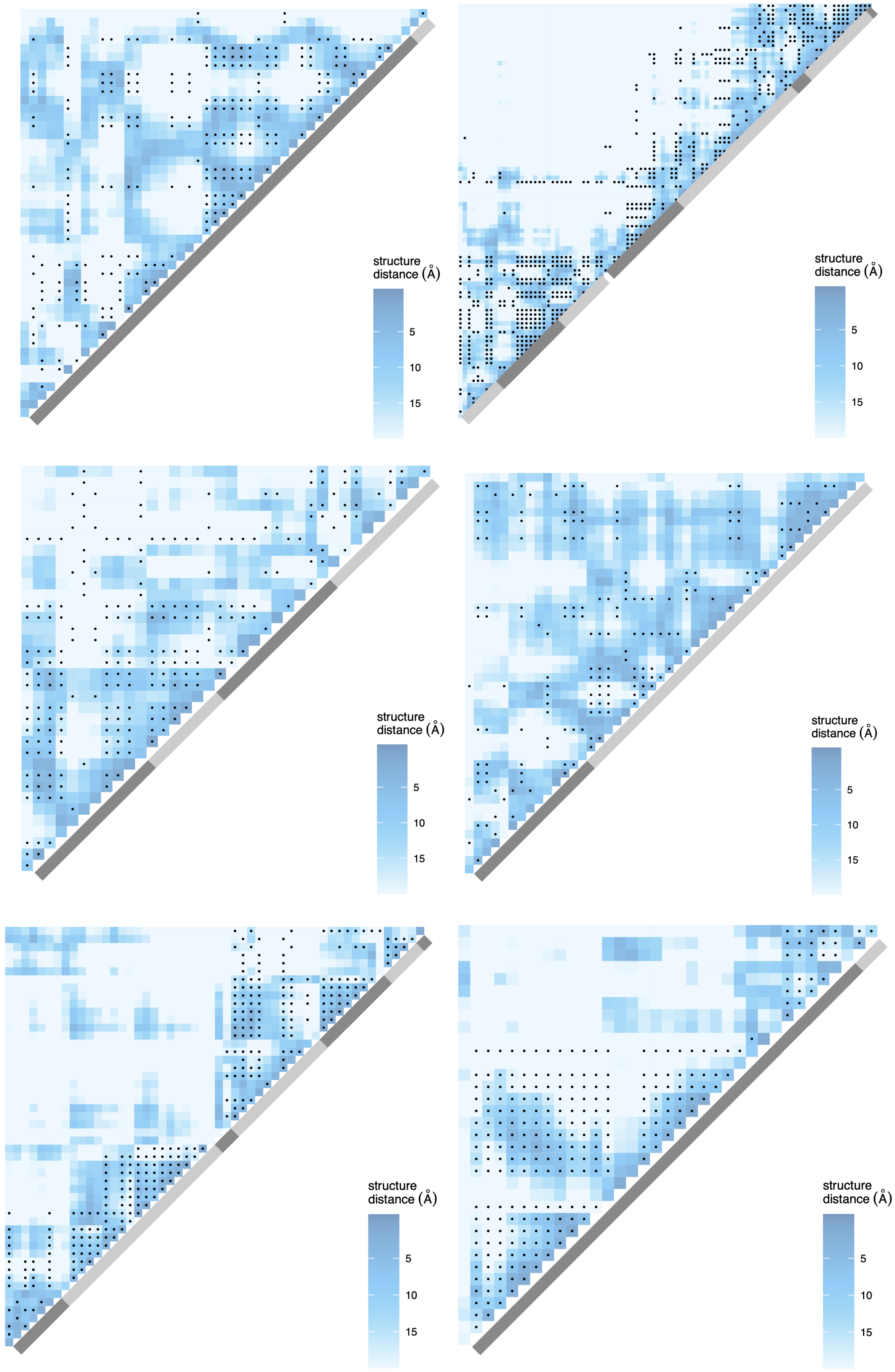
structure distance (Å)
5
10
15
structure distance (Å)
5
10
15
structure distance (Å)
5
10
15
structure distance (Å)
5
10
15
structure distance (Å)
5
10
15
structure distance (Å)
5
10
15

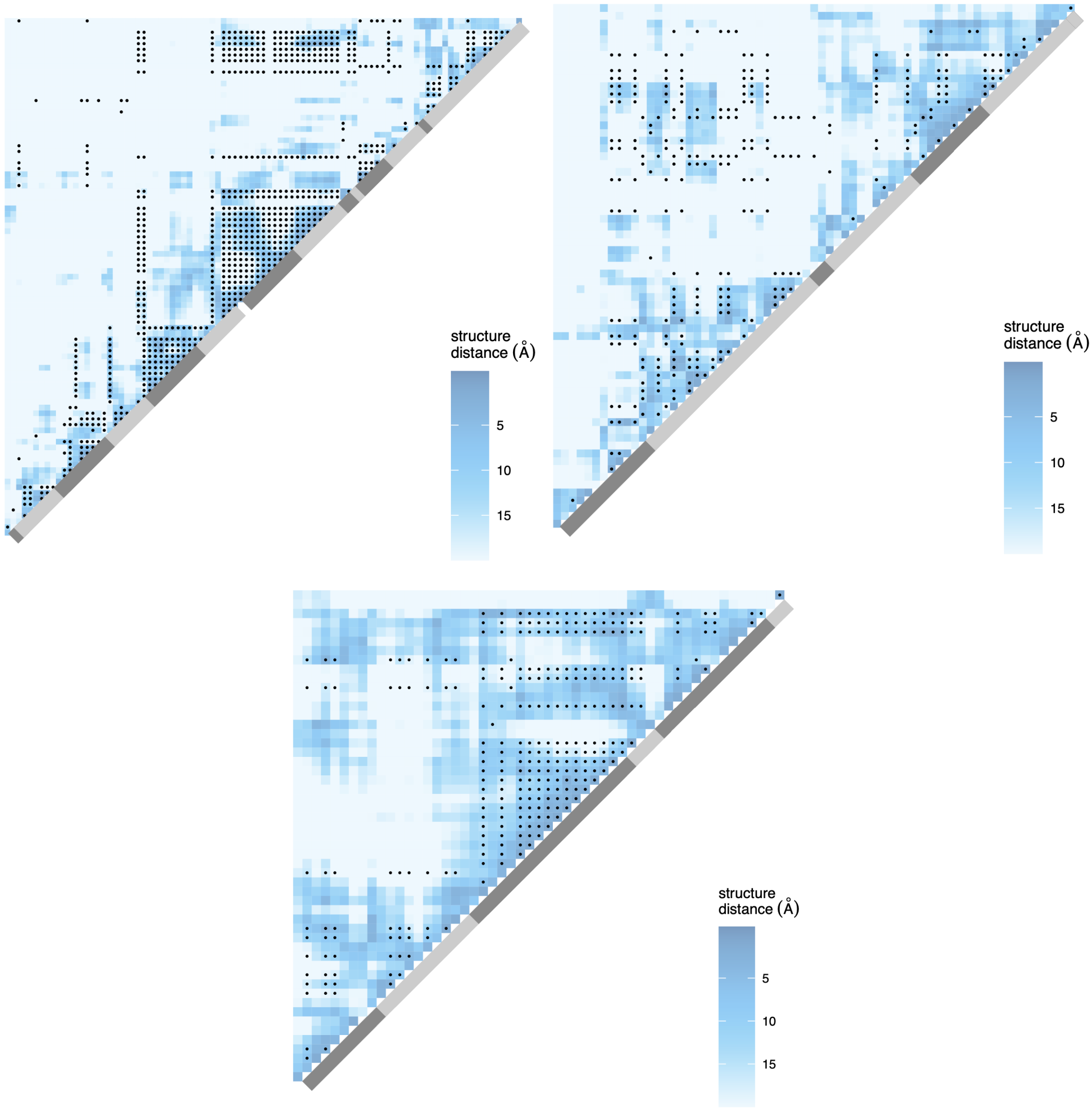


**Supplementary Figure 5. Examples of proteins with LD patterns matching the three-dimensional structure in the RUS population of *S. commune.*** Heatmaps show the physical distance between pairs of sites in the protein structure; only positions carrying biallelic SNPs are shown. Black dots correspond to pairs of sites with high LD (> 0.9 quantile for the gene). Grey regions indicate the exon structure of the genes. (**a**) cog1523 (5Y1B:A); (**b**) cog2779 (1SXJ:B); (**c**) cog3052 (4U9V:B); (**d**) cog5375 (1RGI:G); (**e**), cog5725 (1TA3:B); (**f**) cog14338 (6J3E:A); (**g**) cog18092 (4QJY:A); (**h**) cog7878 (4TYW:A); (**i**) cog18561 (1KSG:A). LD statistics and p-values for each gene are listed in Supplementary Table 1.

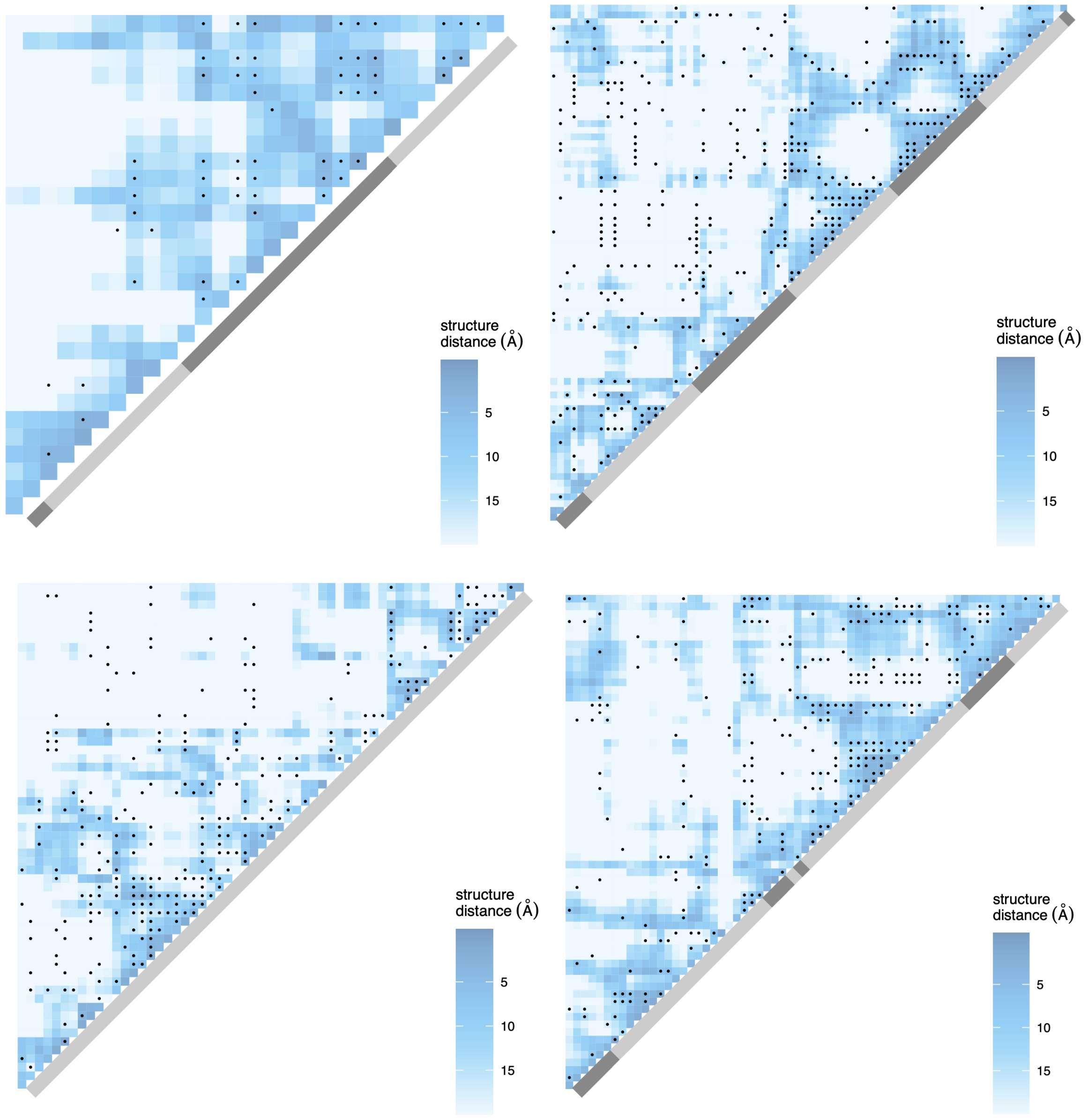


**Supplementary Figure 6. Examples of proteins with LD patterns matching the three-dimensional structure in the USA population of *S. commune*.** Heatmaps show the physical distance between pairs of sites in the protein structure; only positions carrying biallelic SNPs are shown. Black dots correspond to pairs of sites with high LD (> 0.9 quantile for the gene). Grey regions indicate the exon structure of the genes. (**a**) cog1536 (6AHR:E); (**b**) cog5725 (1TA3:B); (**c**) cog8253 (6F87:A); (**d**) cog9241 (1YCD:A). LD statistics and p-values for each gene are listed in Supplementary Table 1.

scaffold4:3097116-3098822

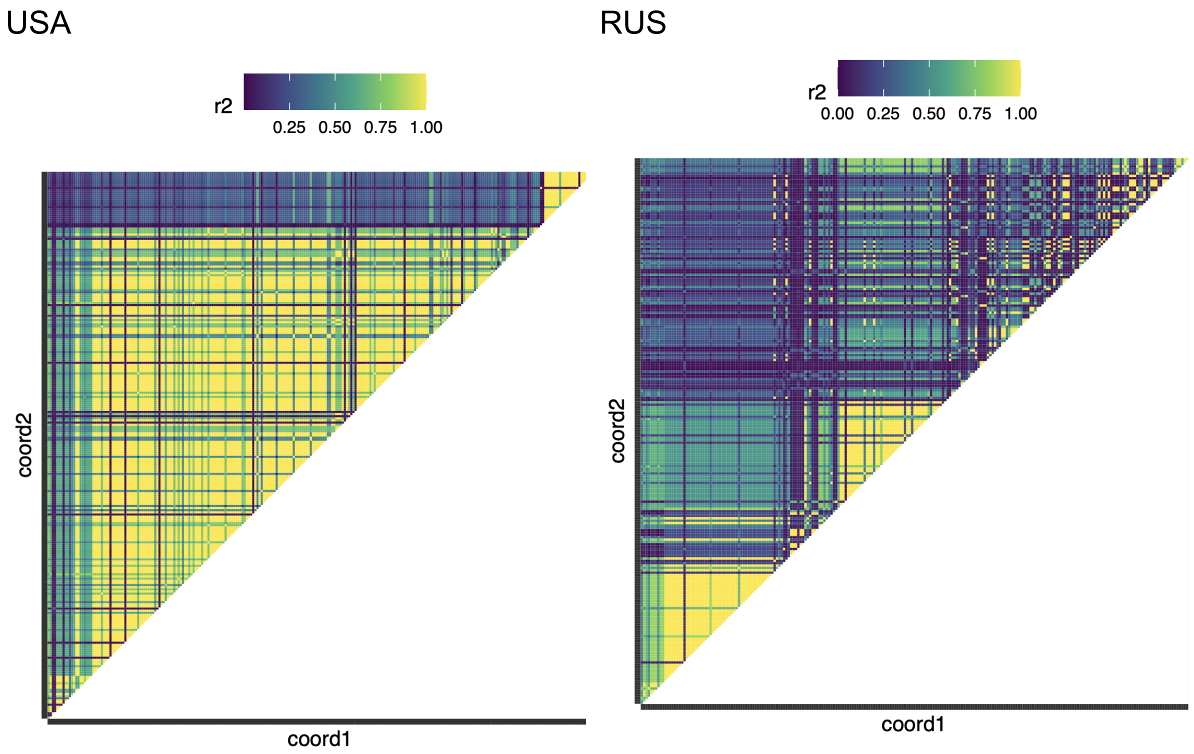


scaffold2:2017262-2019256

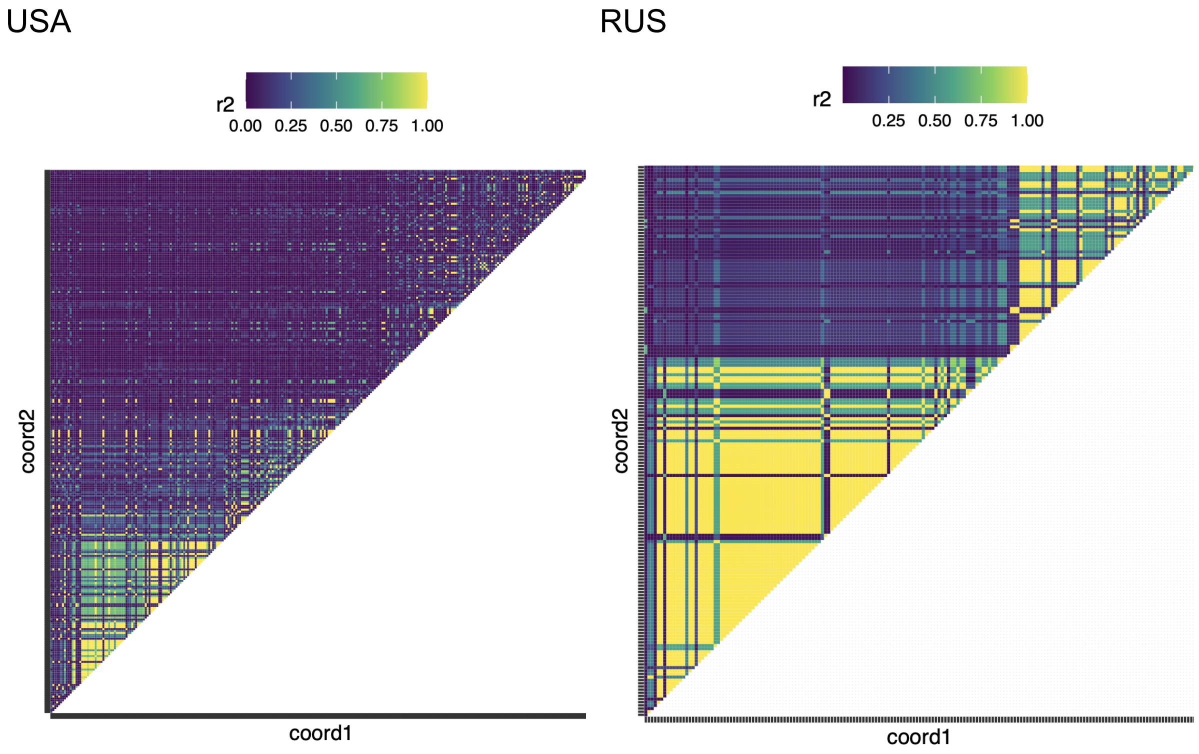

scaffold8:1921168-1924186

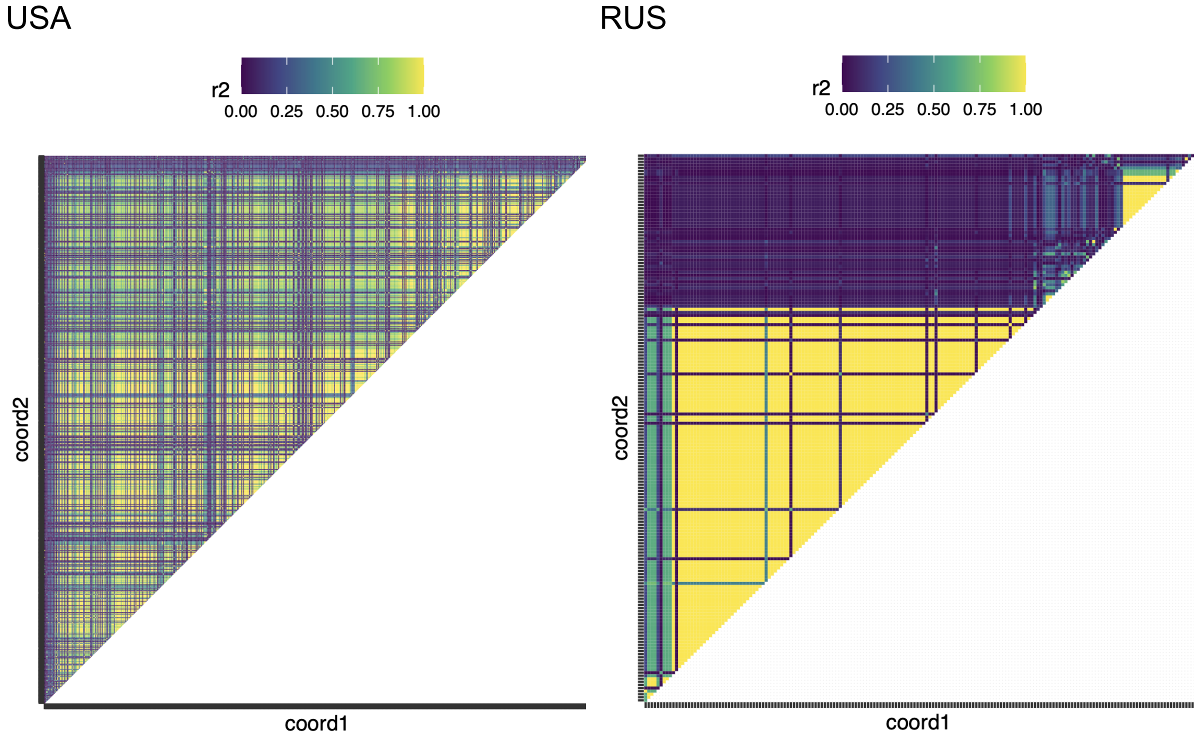


**Supplementary Figure 7. Examples of haploblocks in two populations of *S. commune*.**

The heatmaps show LD between polymorphic SNPs in the same genomic regions in the USA and RUS populations of *S. commune*. Only biallelic polymorphic sites with minor allele frequency > 1 are shown, the number of such sites can differ between populations.

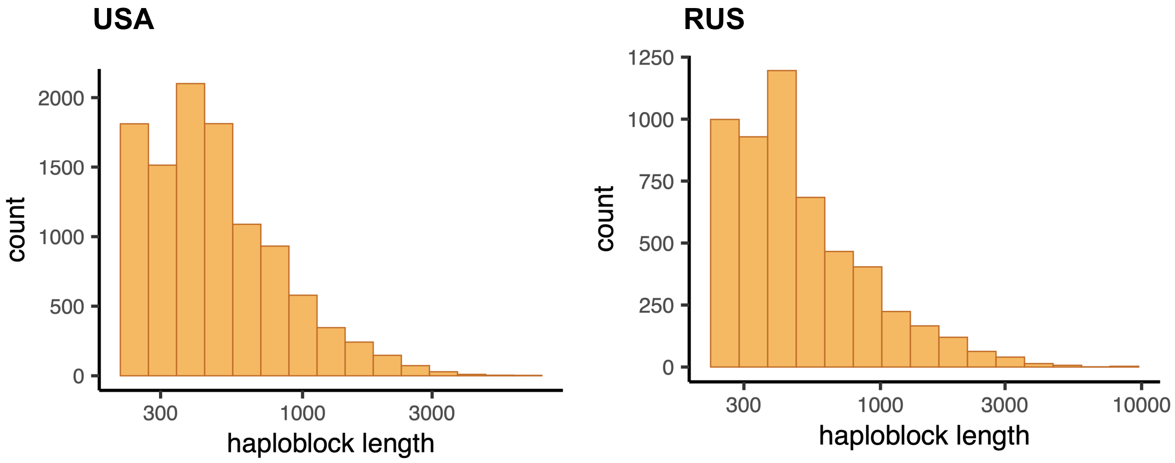


**Supplementary Figure 8. Distribution of haploblock lengths in the two populations of *S. commune*.**

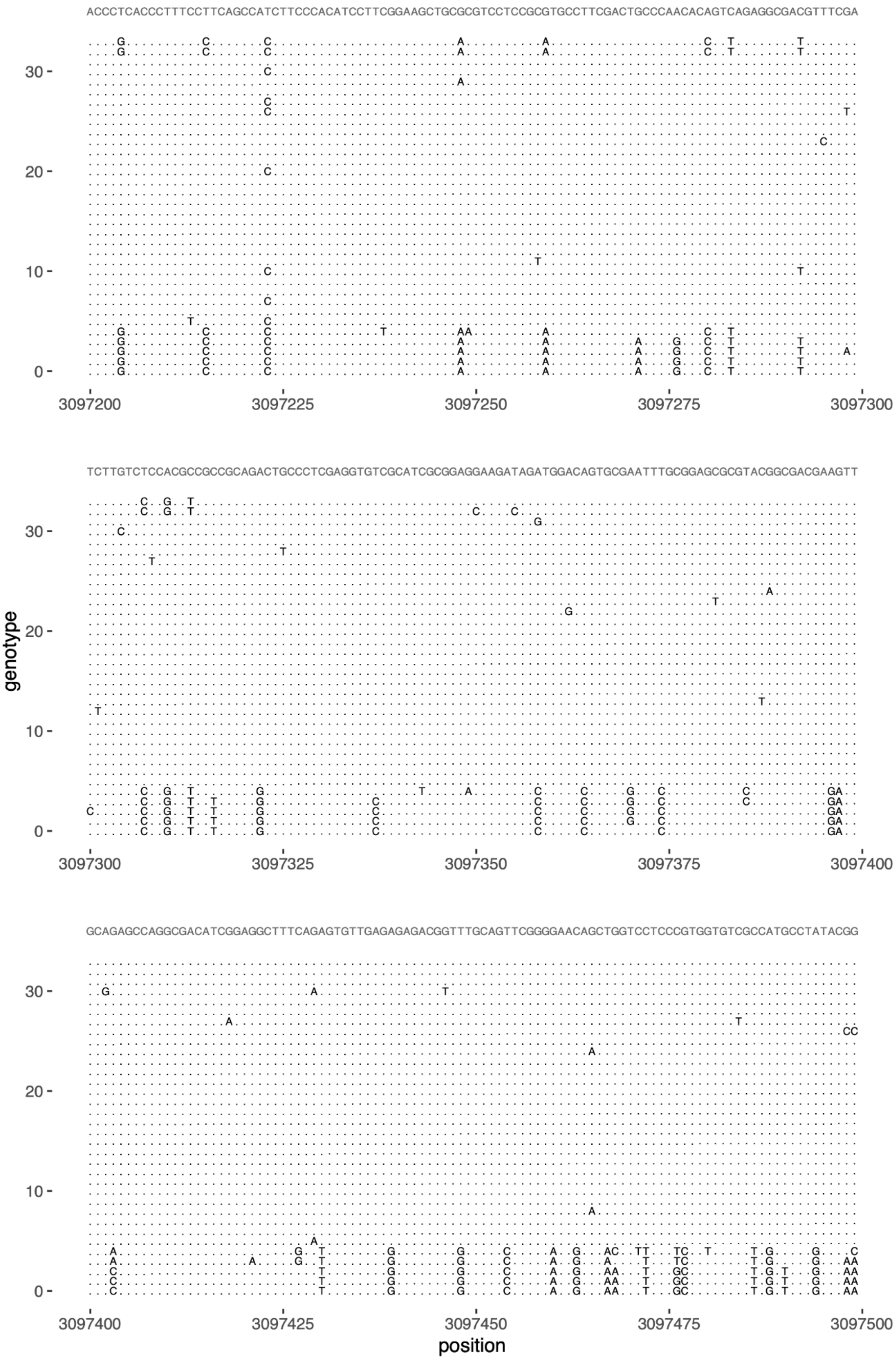


**Supplementary Figure 9. Example of the *S. commune* alignment within a haploblock.** Region 3097200-3097500 of scaffold 4 in the USA population of *S. commune* is shown. The top line shows the consensus sequence based on 34 genotypes; dot indicates match with the consensus.

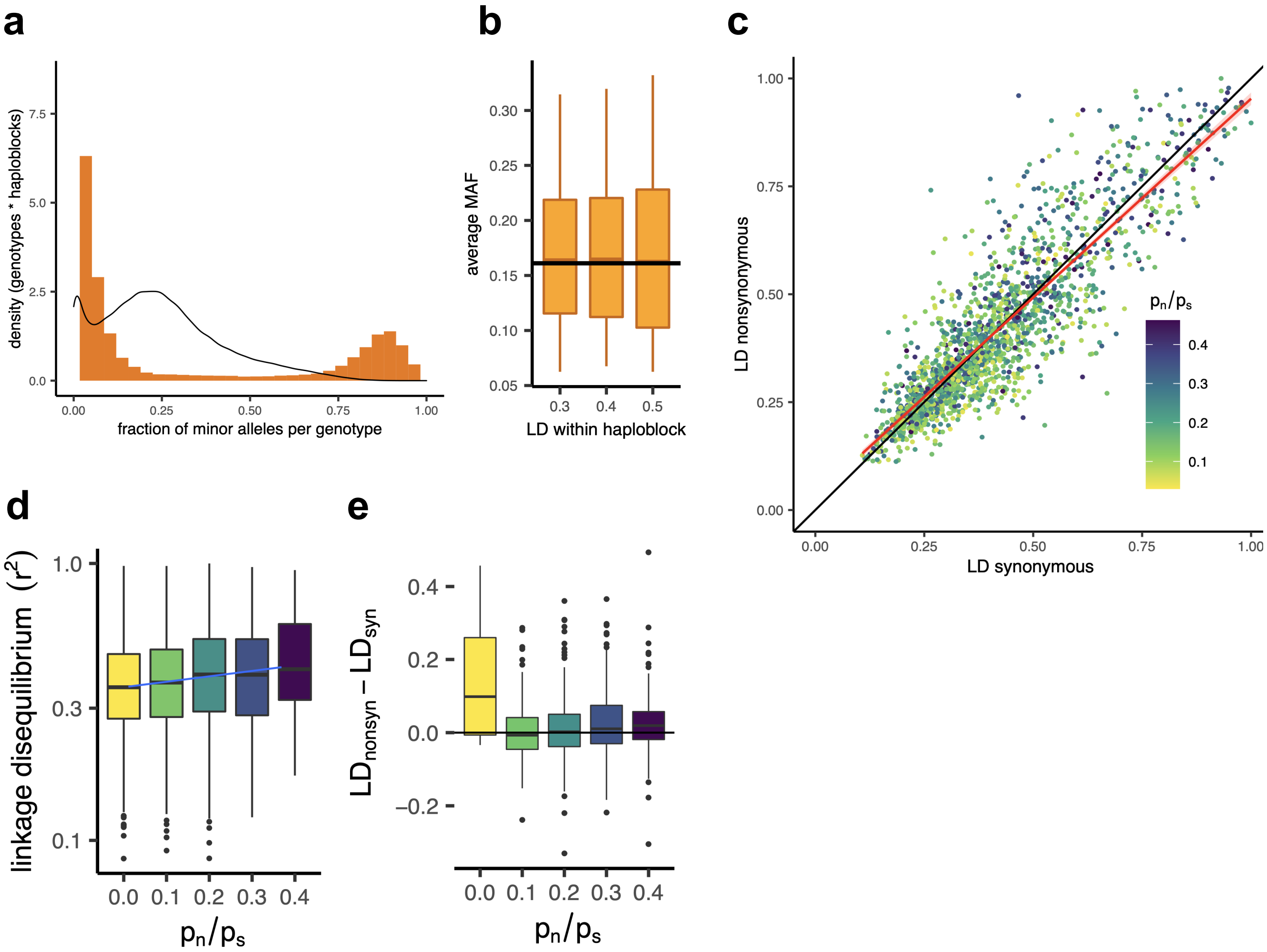


**Supplementary Figure 10. Patterns of linkage disequilibrium in the RUS population of *S. commune*.**

(**a**) Bimodal distribution of the fraction of polymorphic sites carrying minor alleles per genome within the haploblocks. Each count corresponds to a genotype within a haploblock. Black line shows the background distribution of minor alleles in the non-haploblock regions. (**b**) The increased average minor allele frequency within haploblocks as compared to the non-haploblock regions (dashed line, t-test p-value < 2e-16). (**c**) LD between nonsynonymous and synonymous SNPs within single genes. Each dot represents an individual gene. Linear regression of $LD_{nonsyn}$ over $LD_{nsyn}$ is shown as the red line. To control for the gene length, only SNPs within 300 bp from each other were analyzed. Genes with fewer than 100 such pairs of SNPs were excluded. (**d,e**) The positive correlation between $p_n/p_s$ of the gene and its average LD (Spearman correlation p-value = 4e-16) (**d**) or the difference between $LD_{nonsyn}$ and $LD_{syn}$ (Spearman correlation p-value = 2e-5) (**e**).

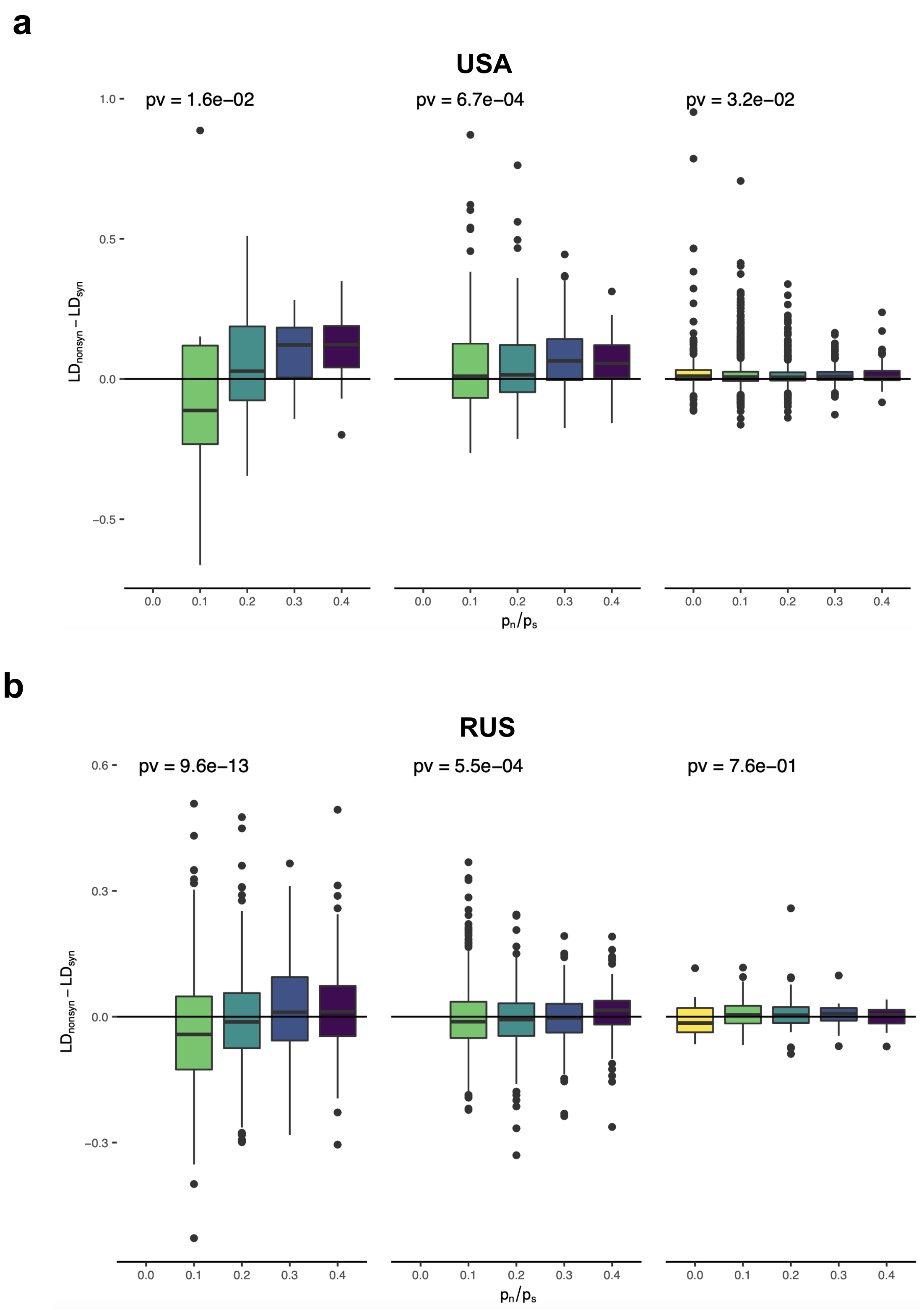


**Supplementary Figure 11. Comparison of $LD_{nonsyn}$ and $LD_{syn}$ in the genes of *S. commune*.** (**a**) The USA population, (**b**) the RUS population. The genes are stratified by their average LD (the panels) and by the $p_n/p_s$. Only pairs of SNPs within 300 bp from each other are analyzed; genes with less than 100 such pairs of nonsynonymous or synonymous SNPs are excluded. Spearman correlation p-values are shown.

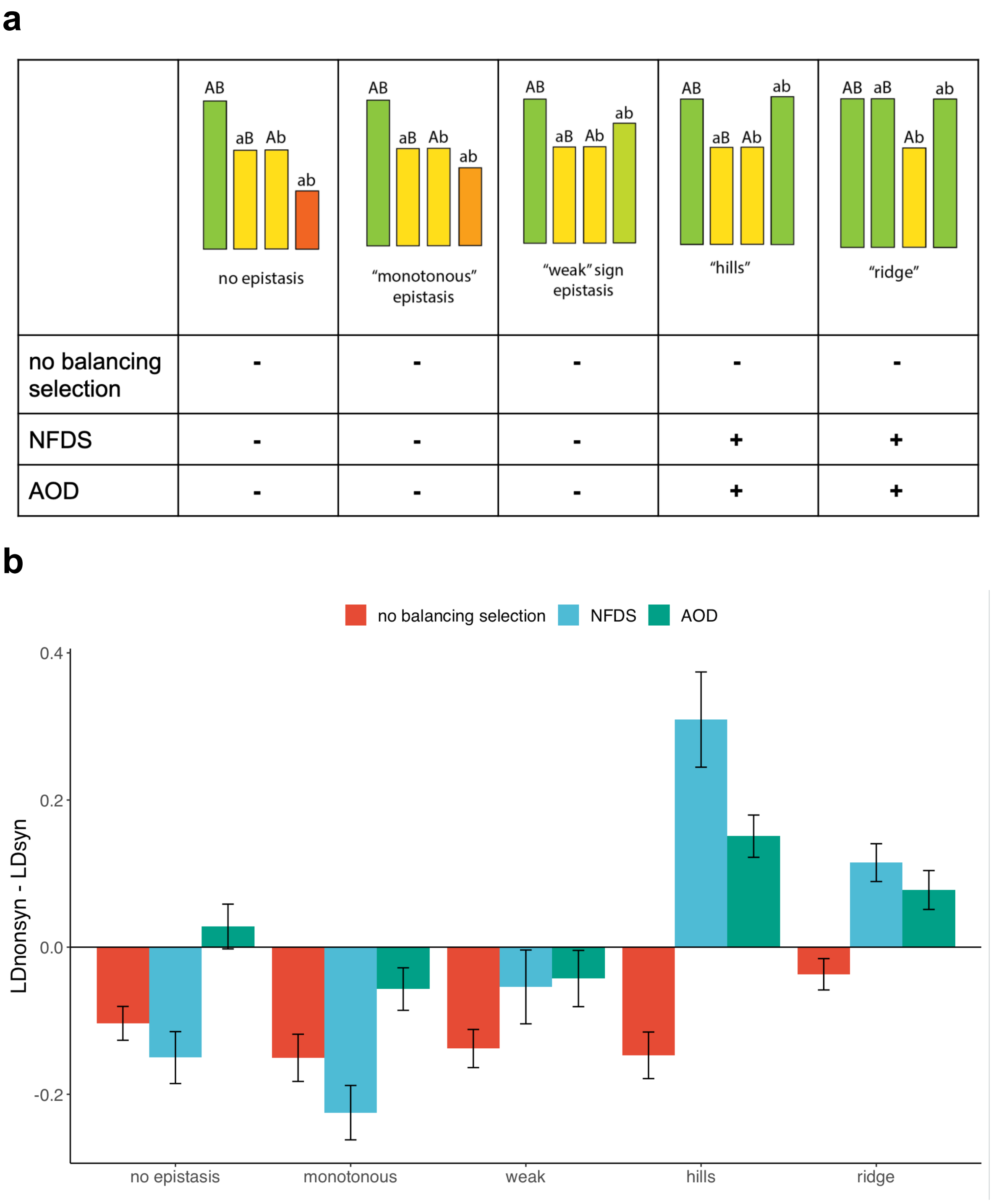


**Supplementary Figure 12. The difference between $LD_{nonsyn}$ and $LD_{syn}$ under pairwise epistasis and balancing selection.** (**a**) The excess of $LD_{nonsyn}$ over $LD_{syn}$ under different models of epistasis between two deleterious mutations A → a and B → b without balancing selection and in the presence of negative frequency-dependent selection (NFDS) or associate overdominance (AOD) acting in the linked sites. The height of columns shows fitness of the corresponding genotypes. (+) indicate simulations where the excess of $LD_{nonsyn}$ is reproduced. (**b**) The difference between $LD_{nonsyn}$ and $LD_{syn}$ in the simulations.

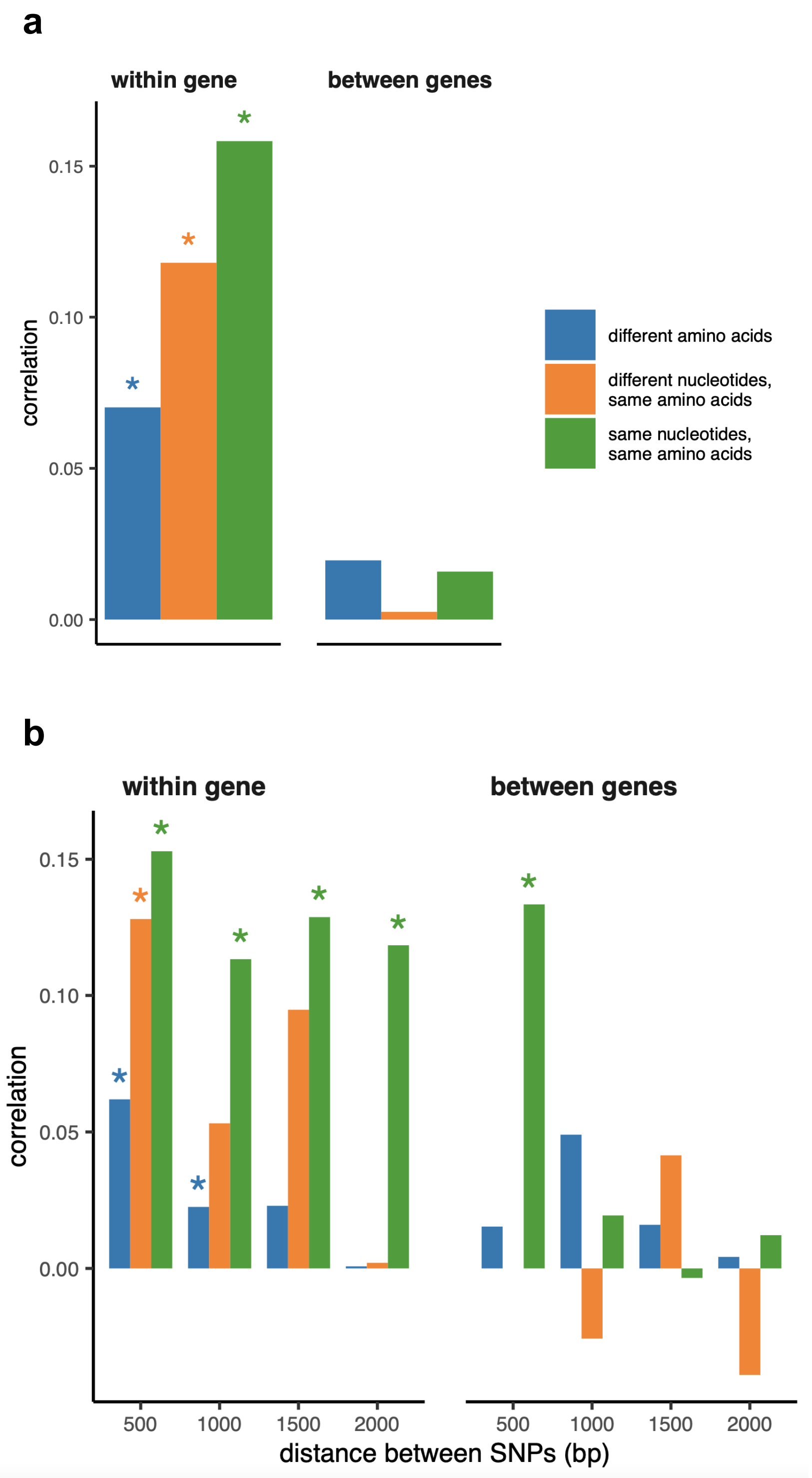


**Supplementary Figure 13. Association of LD values between pairs of shared nonsynonymous SNPs encoding the same amino acids in the two *S. commune* populations.** (**a**) All pairs of SNPs pooled together. Pair of SNPs is considered to carry different alleles if at least one allele differs in at least one site. (**b**) Pairs of SNPs stratified by distance between them. Asterisks indicate Spearman correlation p-values < 0.01.

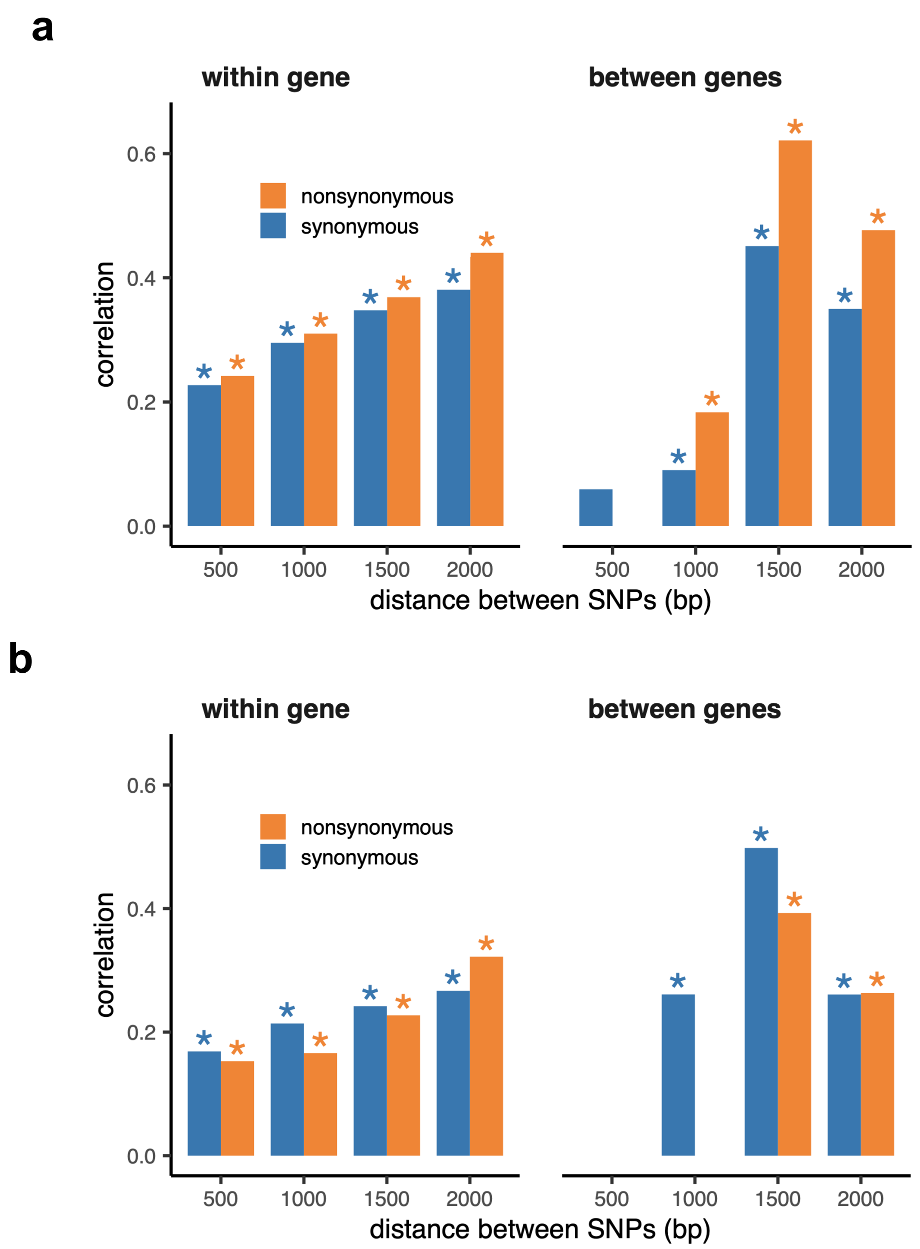


**Supplementary Figure 14. Association of LD values between pairs of shared SNPs within haploblocks in the two *S. commune* populations.** (**a**) Pairs of SNPs with the same major and minor alleles in both sites, (**b**) pairs of SNPs differing by at least one allele. Asterisks indicate Spearman correlation p-values < 0.001.

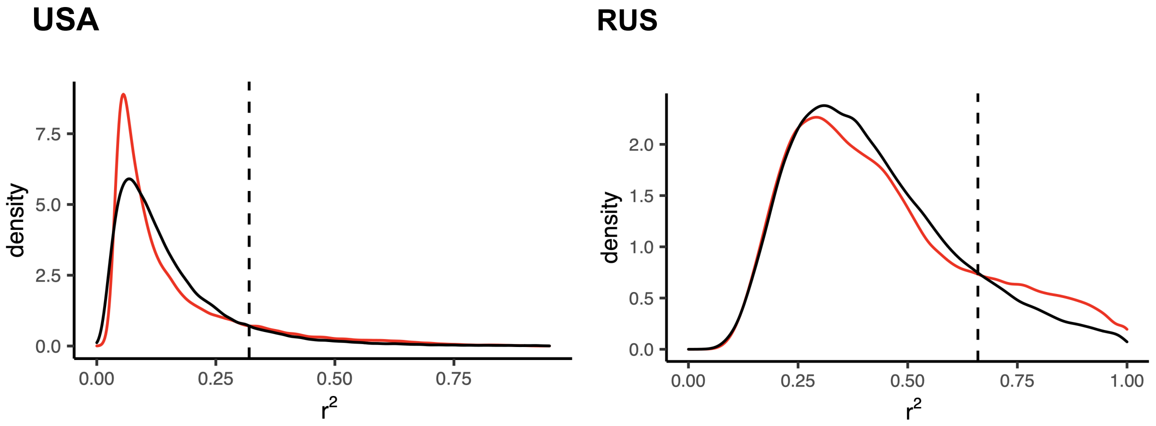


**Supplementary Figure 15. Criteria for haploblocks in *S. commune*.** Red lines show the distribution of LD ($r^2$) in windows of 250 nucleotides in two populations. Black line corresponds to the lognormal distribution with the same mean and variance. The windows with LD higher than the threshold value defined as the intersection point of the two lines (dashed) are attributed to haploblocks.
